# Intrinsic ecological dynamics drive biodiversity turnover in model metacommunities

**DOI:** 10.1101/2020.05.22.110262

**Authors:** Jacob D. O’Sullivan, J. Christopher D. Terry, Axel G. Rossberg

## Abstract

Turnover of species composition through time is frequently observed in ecosystems. It is often interpreted as indicating the impact of changes in the environment. Continuous turnover due solely to ecological dynamics—species interactions and dispersal—is also known to be theoretically possible, however the prevalence of such autonomous turnover in natural communities remains unclear. Here we demonstrate that observed patterns of compositional turnover and other important macroecological phenomena can be reproduced in large spatially explicit model ecosystems, without external forcing such as environmental change or the invasion of new species into the model. These results imply that the potential role of autonomous turnover as a widespread and important natural process is underappreciated, challenging assumptions implicit in many observation and management tools. Quantifying the baseline level of compositional change would greatly improve ecological status assessments.

**One Sentence Summary:** Biodiversity change previously attributed to external drivers is explainable as resulting from intrinsic ecosystem dynamics.

## INTRODUCTION

Change in species composition observed in a single locality through time, called community turnover, is observed to occur in most ecosystems at a faster rate than is explainable by random drift^1,2^. Climate change and other anthropogenic pressures are known to contribute to community turnover^3–6^ and there is evidence to suggest that turnover is accelerating in some biomes^7^. The extent to which processes intrinsic to ecosystems contribute to turnover, however, remains poorly understood^8^.

Previous theoretical^9,10^ and experimental studies^11^ have shown how specific motifs in competitive ecological networks can lead to population abundances which do not arrive at fixed points. Instead, such systems can manifest persistent dynamics which we refer to here as ‘autonomous’ since they do not depend on variation in the external environment or other extrinsic drivers. When these population fluctuations are strong, changes in the abundances of species can be dramatic and even drive species locally extinct; if an excluded species retains occupancy in adjacent patches^10^, it may re-colonise at some future time. We refer to as ‘autonomous turnover’ local compositional changes involving colonisation-extinction processes or significant restructuring of relative abundances, driven by autonomous population dynamics.

Understanding the expected amount of autonomous turnover in natural systems is important if change in the composition of ecological communities is to be interpreted as indicative of community stress^12,13^. If strong temporal community turnover was a natural phenomenon that can arise independently of changes in the abiotic environment, then observed shifts in the composition of ecological communities would not on their own carry the fingerprint of external pressures.

Limitations in the availability of historical turnover rates before the onset of widespread anthropogenic impacts pose considerable challenges when trying to establish the natural baseline of turnover. Nevertheless, emergent patterns in species-time-area relationships^14,15^ suggest an underlying consistency accessible through modelling.

Antagonistic interactions between predators and prey have been shown in both theory and experiment to lead to persistent population oscillations in the absence of external variation^16,17^. It has also long been established that models of competitive communities can generate any type of dynamical behaviour, including persistent chaotic cycles^18–20^. However, these cyclic processes are different from and have not usually been associated with observations of acyclic compositional turnover^1,2^. An important distinction between these processes lies in the role of space. While cyclic forms of community dynamics can lead to characterisic spacial structures^20,21^, cyclic dynamics do principle not require space. Acyclic turnover, on the other hand, manifestly involves colonisation by species from surrounding patches.

Here we ask: can community dynamics enabled by spatial structure account for the observed macroecological patterns in population turnover? We address this question drawing on recent advances in the theory of spatially extended ecological communities, so called metacommunities^22^, using a population-dynamical simulation model with explicit spatial and environmental structure^23^ that has previously been shown to reproduce fundamental *spatial* biodiversity patterns. Here we build upon this work by exploring the spatio-*temporal* patterns that emerge in metacommunity models. As shown below, these arise when expanding the spatial and taxonomic scale of simulations beyond those studied previously.

## RESULTS

### Metacommunity model and asymptotic community assembly

We built a large set of model metacommunities (detailed in full in Methods) describing competitive dynamics within a single guild of species across a landscape. Each metacommunity consisted of a set of patches, or local communities, randomly placed in a square arena and linked by a spatial network. The dynamics of each population are governed by three processes: inter- and intraspecific interactions, heterogeneous responses to the environment and dispersal between adjacent patches (Fig. 1). Competition coefficients between species are drawn at random and the population dynamics within each patch are described by a Lotka-Volterra competition model. We control the level of environmental heterogeneity across the network directly by generating an intrinsic growth rate for each species at each patch from a random, spatially correlated distribution. To ensure any turnover is purely autonomous, we keep the environment fixed throughout simulations. Dispersal between neighbouring patches declines exponentially with distance between sites. This formulation allows precise and independent control of key properties of the metacommunity–the number of patches, the characteristic dispersal length and the heterogeneity of the environment.

**Figure 1.**
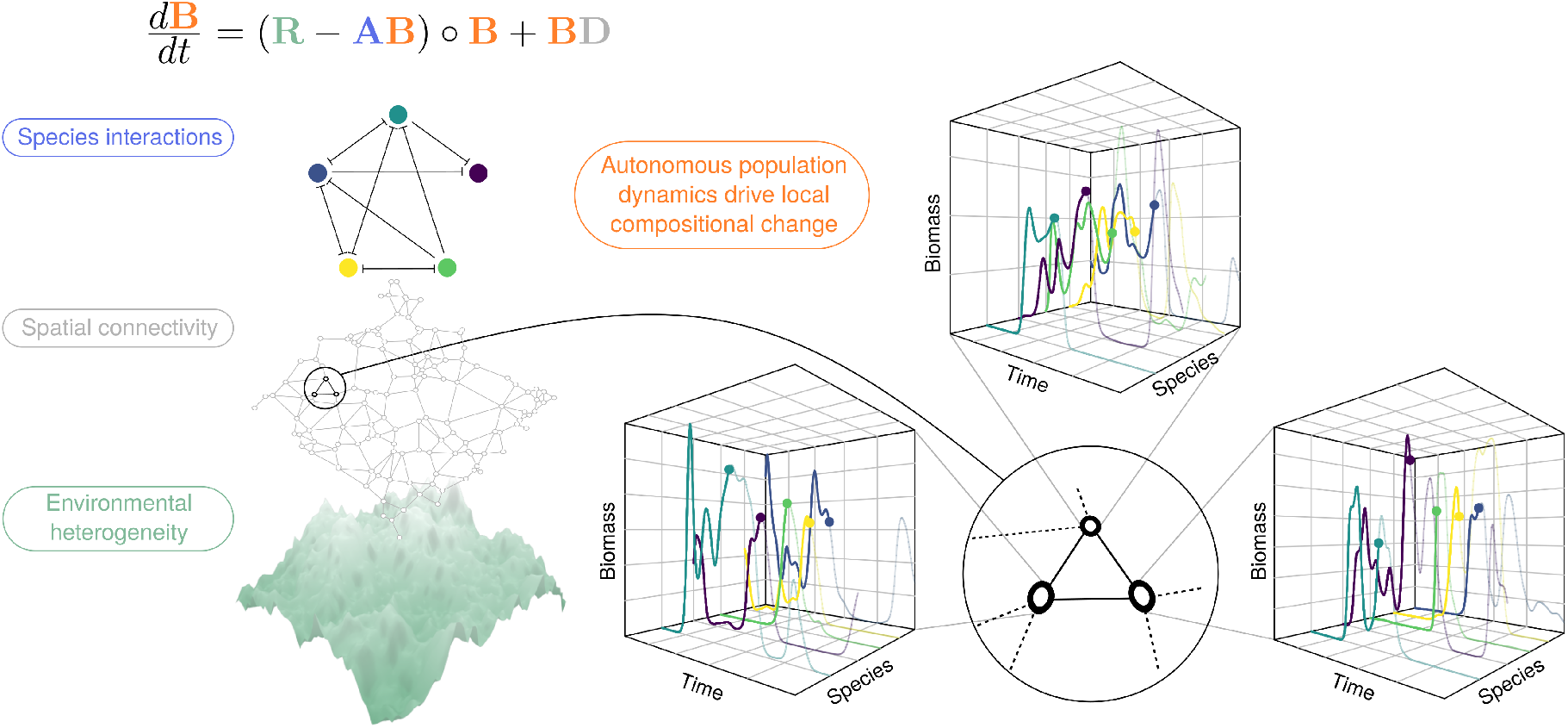
Structure of the Lotka-Volterra metacommunity model and emergence of autonomous population dynamics. *Environmental heterogeneity* is modelled using a spatially autocorrelated Gaussian random field. A random spatial network defines the *spatial connectivity* of the landscape. The network of *species interactions* is modelled by sampling competition coefficients at random (perpendicular bars indicate recipients of a deleterious competitive impact). The resulting dynamics of local population biomasses, given by colour-coded equation, are numerically simulated. For large metacommunities, local populations exhibit persistent dynamics despite absence of external drivers. In the 3D boxes, typical simulated biomass dynamics of dominating species are plotted on linear axes over 2500 unit times. The graphs illustrate the complexity of the autonomous dynamics and the propensity for compositional change (local extinction and colonisation).

To populate the model metacommunities, we iteratively introduced species with randomly generated intrinsic growth rates and interspecific interaction coefficients. Between successive invasions we simulated the model dynamics, and removed any species whose abundance fell below a threshold across the whole network. Through this assembly process both the average *local diversity*, the number of species coexisting in a given patch, and the *regional diversity*, the total number of species in the metacommunity, eventually saturate and then fluctuate around an equilibrium value—any introduction of a new species then leads on average to the extinction of one other species (Fig. S1). As previously shown^23^, these metacommunities have reached a state of ‘ecological structural instability’^24^, as a result of which species richness is intrinsically regulated. In these metacommunities we then studied the phenomenology of autonomous community turnover *in the absence of regional invasions or abiotic change*.

The structural stability of a system refers to its capacity to sustain changes in parameters without undergoing qualitative changes in dynamical behaviour^25^. In ecological communities, for which abundances are all strictly non-negative, an important qualitative change occurs when species are driven extinct. As such, ecological structural stability is taken to describe in particular the capacity of a community to *persist* (all constituent species have abundances greater than zero) in the face of small biotic or aboitic perturbations^24,26–30^. Ecological structural instability has been shown to play a critical role in the regulation of biodiversity, setting hard limits to the number of species that can coexist^24^, a mechanism found to operate at both local and metacommunity scales^23^. Empirical observation of many of the emergent phenomena associated with ecological structural instability provides strong indirect evidence for the prevalence of structural instability in nature^23,31^. Our understanding of the impact of structurally unstable diversity regulation on temporal community-level properties, however, remains incomplete^32^.

In our metacommunity model, local community dynamics and therefore local limits on species richness depend on a combination of abiotic and biotic *filtering* (non-uniform responses of species to local conditions)^33–35^ and immigration from adjacent patches, generating so called *mass effects* in the local community^36–38^. Abiotic filtering occurs via the spatial variation of intrinsic growth rates *R_ix_* and biotic filtering via interspecific competition encoded in the interaction coefficients *A_ij_*. Intrinsic growth rates *R_ix_* are sampled from spatially correlated normal distributions with mean 1, autocorrelation length *φ* and variance *σ*^2^ (Fig. S2). For simplicity, and since predator-prey dynamics are known to generate oscillations^39^ through mechanisms distinct from those we report here, we restrict our analysis to competitive communities for which all ecological interactions are antagonistic. The off-diagonal elements of the interaction matrix **A**, describing how one species *i* affects another species *j*, are sampled independently from a discrete distribution, such that the interaction strength *A_ij_* is set to a constant value in the range 0 to 1 (in most cases 0.5) with fixed probability (connectance, in most cases 0.5) and otherwise set to zero. Intraspecific competition coefficients *A_ii_* are set to 1 for all species.

Dispersal is modelled via a spatial connectivity matrix with elements *D_xy_*. The topology of the model metacommunity, expressed through **D**, is generated by sampling the spatial coordinates of *N* patches from a uniform distribution 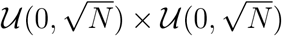, i.e., an area of size *N*. Thus, under variation of the number of patches, the inter-patch distances remain fixed on average. Spatial connectivity is defined by linking these patches through a Gabriel graph^40^, a planar graph generated by an algorithm that, on average, links each local community to four close neighbours^41^. Avoidance of direct long-distance dispersal and the sparsity of the resulting dispersal matrix permit the use of efficient numerical methods. The exponential dispersal kernel defining *D_xy_* is tuned by the dispersal length *ℓ*, which is fixed for all species.

The dynamics of local population biomasses *B_ix_ = B_ix_*(*t*) are modelled using a system of spatially coupled Lotka-Volterra (LV) equations that, in matrix notation, takes the form^23^

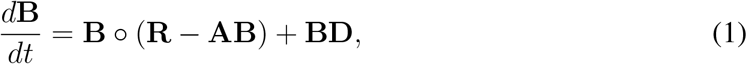

with ◯ denoting element-wise multiplication. Hereafter this formalism is referred to as the Lotka-Volterra Metacommunity Model (LVMCM). Further technical details are provided in Methods and the Supplementary material.

In order to numerically probe the impact of *ℓ, φ* and *σ*^2^ on the emergent temporal dynamics, we initially fixed *N* = 64 and varied each parameter through multiple orders of magnitude (Fig. S3). In order to obtain a full characterisation autonomous turnover in the computationally accessible spatial range (*N* ≤ 256), we then selected a parameter combination found to generate substantial fluctuations for further analysis. Thereafter we assembled metacommunities of 8 to 256 patches (Fig. 2A) to regional saturation (with 10-fold replication) and generated community time series of 10^4^ unit times from which the phenomenology of autonomous turnover could be explored in detail. We found no evidence to suggest that the phenomenology described below depends on this specific parameter combination. While future results may confirm or refute this, autonomous turnover arises over a wide range of parameters (Fig. S3) and as such the phenomenon is reasonably robust.

**Figure 2.**
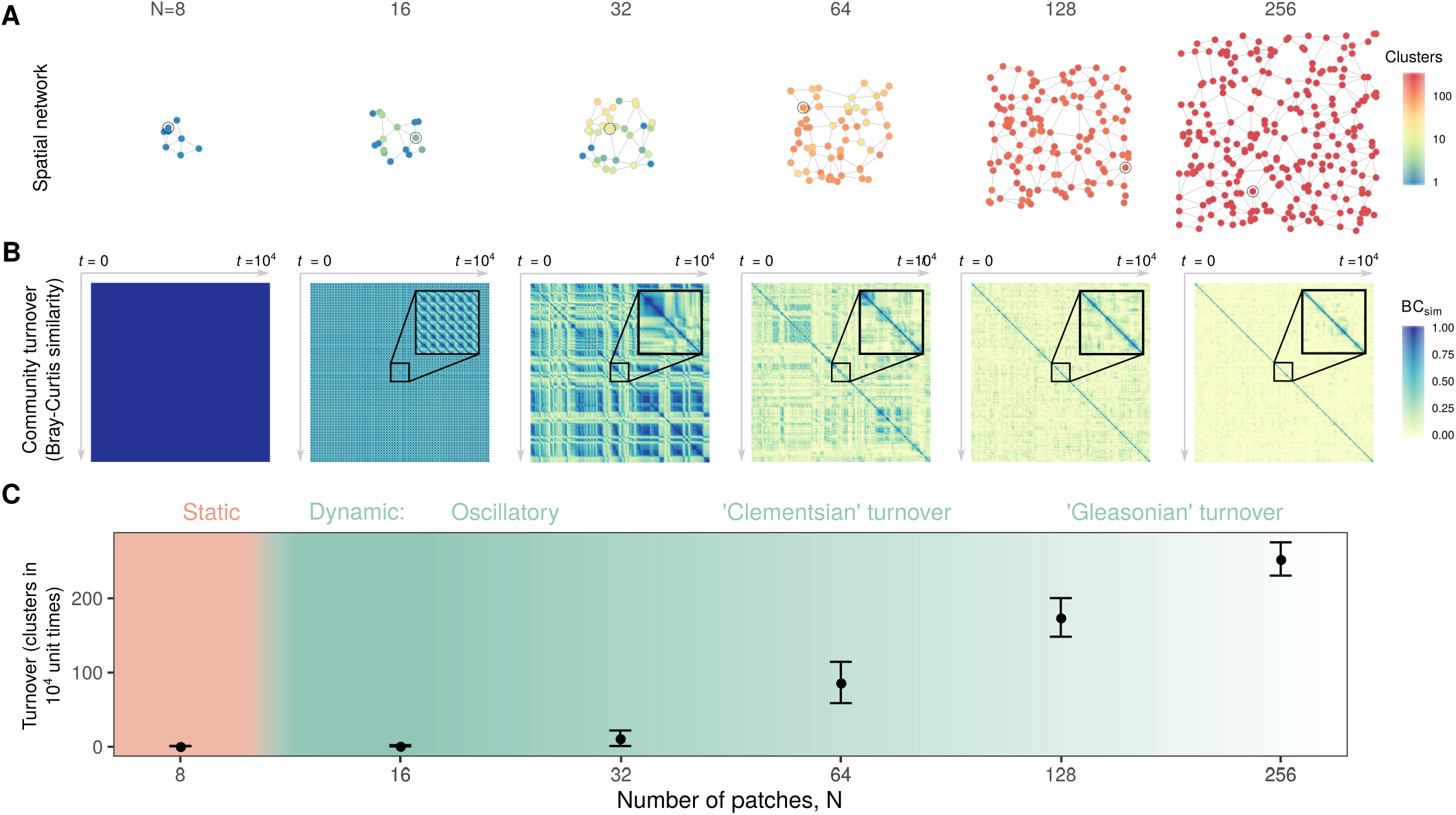
Autonomous turnover in model metacommunities. **A**: Typical model metacommunities: a spatial network with nodes representing local communities (or patches) and edges, channels of dispersal. Patch colour represents the number of clusters in local community state space detected over 10^4^ unit times using hierarchical clustering of the Bray-Curtis (BC) similarity matrix, Fig. S5. **B**: Colour coded matrices of pairwise temporal BC similarity corresponding to the circled patches in **A**. Insets represent 10^2^ unit times. For small networks (*N* = 8) local compositions converge to static fixed points. As metacommunity extent increases, however, persistent dynamics emerge. Initially this autonomous turnover is oscillatory in nature with communities fluctuating between small numbers of states which can be grouped into clusters (16≤N ≤32). Intermediate metacommunities (32 ≤N ≤64) manifest ‘Clementsian’ temporal turnover, characterized by sharp transitions in composition, implying species turn over in cohorts. Large metacommunities (N ≥ 128) turn over continuously, implying ‘Gleasonian’ assembly dynamics in which species’ temporal occupancies are independent. **C**: The mean number of local compositional clusters detected for metacommunities of various numbers of patches N. While the transition from static to dynamic community composition at the local scale is sharp (see text), nonuniform turnover *within* metacommunities (**A**) blurs the transition at the regional scale. *A_ij_* = 0.5 with probability 0.5, *φ* = 10, *σ*^2^ = 0.01, *ℓ* = 0.5.

### Autonomous turnover in model metacommunities

For small (*N* ≤ 8) metacommunities assembled to saturation of regional diversity, populations attain equilibria, i.e. converge to fixed points, implying the absence of autonomous turnover^23^. With increasing metacommunity size *N*, however, we observe the emergence of persistent population dynamics (Fig. S4, https://vimeo.com/379033867) that can produce substantial turnover in local community composition. This autonomous turnover can be represented through Bray-Curtis^42^ (BC) similarity matrices comparing local community composition through time (Fig. 2B), and quantified by the number of compositional clusters detected in such matrices using hierarchical cluster analysis (Fig. 2A and C).

At intermediate spatial scales (Fig. 2, 16 ≤ *N* ≤ 32) we often find oscillatory dynamics, which can be perfectly periodic or slightly irregular. With increasing oscillation amplitude, these lead to persistent turnover dynamics where local communities repeatedly fluctuate between a small number of distinct compositional clusters (represented in Fig. 2 by stripes of high pairwise BC similarity spanning large temporal ranges). At even larger scales (*N* ≥ 64) this compositional coherence begins to break down, and for very large metacommunities (*N* ≥ 128) autonomous dynamics drive continuous and unpredictable change in community composition. The number of compositional clusters detected typically varies within a given metacommunity (Fig. 2A node colour), however we find a clear increase in the average number of compositional clusters, i.e. an increase in turnover, with increasing total metacommunity size (Fig. 2C).

Metacommunities in which the boundaries of species ranges along environmental gradients are clumped are termed *Clementsian*, while those for which range limits are independently distributed are denoted *Gleasonian*^43^. We consider the block structure of the temporal dissimilarity matrix at intermediate *N* to represent a form of Clementsian temporal turnover, characterized by sudden significant shifts in community composition. Metacommunity models similar to ours have been found to generate such patterns along spatial gradients^44^, potentially via an analogous mechanism^45^. Large, diverse metacommunities manifest Gleasonian temporal turnover. In such cases, species colonisations and extirpations are largely independent and temporal occupancies predominantly uncorrelated, such that compositional change is continuous, rarely, if ever, reverting to the same state.

### Mechanistic explanation of autonomous turnover

Surprisingly, the onset and increasing complexity of autonomous turnover as system size *N* increases (Fig. 2) can be understood as a consequence of local community dynamics alone. To explain this, we first recall relevant theoretical results for isolated LV communities. Then we demonstrate that, in presence of weak propagule pressure, these results imply local community turnover dynamics, controlled by the richness of potential invaders, that closely mirror the dependence on system size seen in full LV metacommunities.

Application of methods from statistical mechanics to models of large isolated LV communities with random interactions revealed that such models exhibit qualitatively distinct phases^49–51^. If the number of modelled species, *S*, interpreted as species pool size, lies below some threshold value determined by the distribution of interaction strengths (Fig. S6), these models exhibit a unique linearly stable equilibrium (Unique Fixed Point phase, UFP). Some species may go extinct, but the majority persists^51^. When pool size *S* exceeds this threshold, there appear to be no more linearly stable equilibrium configurations. Any community formed by a selection from the *S* species is either unfeasible (there is no equilibrium with all species present), intrinsically linearly unstable, or invadable by at least one of the excluded species. This has been called the multiple attractor (MA) phase^50^. However, the precise nature of dynamics in this MA phase appears to remain unclear.

Population dynamical models with many species have been shown to easily exhibit attractors called stable heteroclinic networks^46^, which are characterized by dynamics in which the system bounces around between several unstable equilibria, each corresponding to a different composition of the extant community, implying indefinite community turnover (Fig. 3, red line). As these attractors are approached, models exhibit increasingly long intermittent phases of slow dynamics, which, when numerically simulated, can give the impression that the system eventually reaches one of several ‘stable’ equilibria. We demonstrate in supplementary text that the MA phase of isolated LV models is in fact characterized by such stable heteroclinic networks (Figs. S7, S8).^52^

If one now adds to such isolated LV models terms representing weak propagule pressure for all *S* species (Eq. S5), dynamically equivalent to mass effects occurring in the full metacommunity model (Eq. 1), then none of the *S* species can entirely go extinct. The weak influx of biomass drives community states away from the unstable equilibria representing coexistence of subsets of the *S* species and the heteroclinic network connecting them (blue line in Fig. 3). Typically, system dynamics then still follow trajectories closely tracking the original heteroclinic networks (Fig. 3), but now without requiring boundless time to transition from the vicinity of one equilibrium to the next.

**Figure 3.**
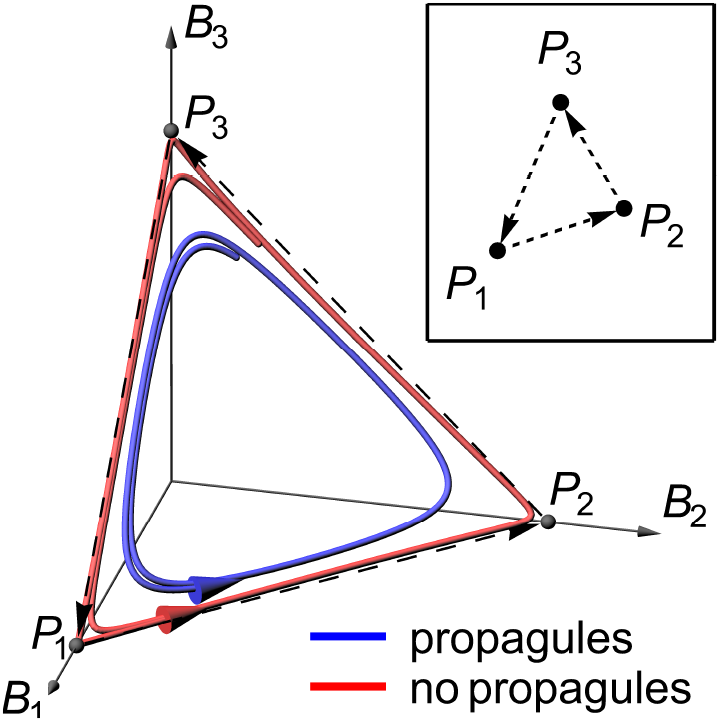
Approximate heteroclinic networks underlie autonomous community turnover. The main panel shows two trajectories in the state space of a community of three hypothetical species (population biomasses *B*_1_, *B*_2_, *B*_3_) that are in non-hierarchical competition with each other, such that no species can competitively exclude both others (a “rock-paper-scissors game”^20^). Without propagule pressure, the system has three unstable equilibrium points (*P*_1_, *P*_2_, *P*_3_) and cycles between these (red curve), coming increasingly close to the equilibria and spending ever more time in the vicinity of each. The corresponding attractor is called a *heteroclinic cycle* (dashed arrows). Under weak extrinsic propagule pressure (blue curve), the three equilibria and the heteroclinic cycle disappear, yet the system closely tracks the original cycle in state space. Such a cycle can be represented as a graph linking the dynamically connected equilibria (inset). With more interacting species, these graphs can become complex “heteroclinic networks”^46–48^ representing complex sequences of species composition during autonomous community turnover.

The nature and complexity of the resulting population dynamics depend on the size and complexity of the underlying heteroclinic network, and both increase with pool size *S*. In simulations (Fig. S9) we find that, as *S* increases, LV models with weak propagule pressure pass through the same sequence of states as we documented for LVMCM metacommunities in Fig. 2: equilibria, oscillatory population dynamics, Clementsian and finally Gleasonian temporal turnover.

Above we introduced the number of clusters detected in Bray-Curtis similarity matrices of fixed time series length as a means of quantifing the approximate number of equilibria visited during local community turnover. As shown in Fig. 4, this number increases in LV models with *S* in a manner strikingly similar to its increase in the LVMCM with the number of species present in the ecological neighbourhood of a given patch. Thus dynamics within a patch are controlled not by *N* directly but rather by neighbourhood species richness which, due to spatial inhomogeneities, varies from patch to patch for metacommunities of a given size *N*. As illustrated in Fig. 4, there is a tendency for neighbourhood richness to be larger in larger metacommunities, leading indirectly to the dependence of metacommunity dynamics on *N* seen in Fig. 2.

**Figure 4.**
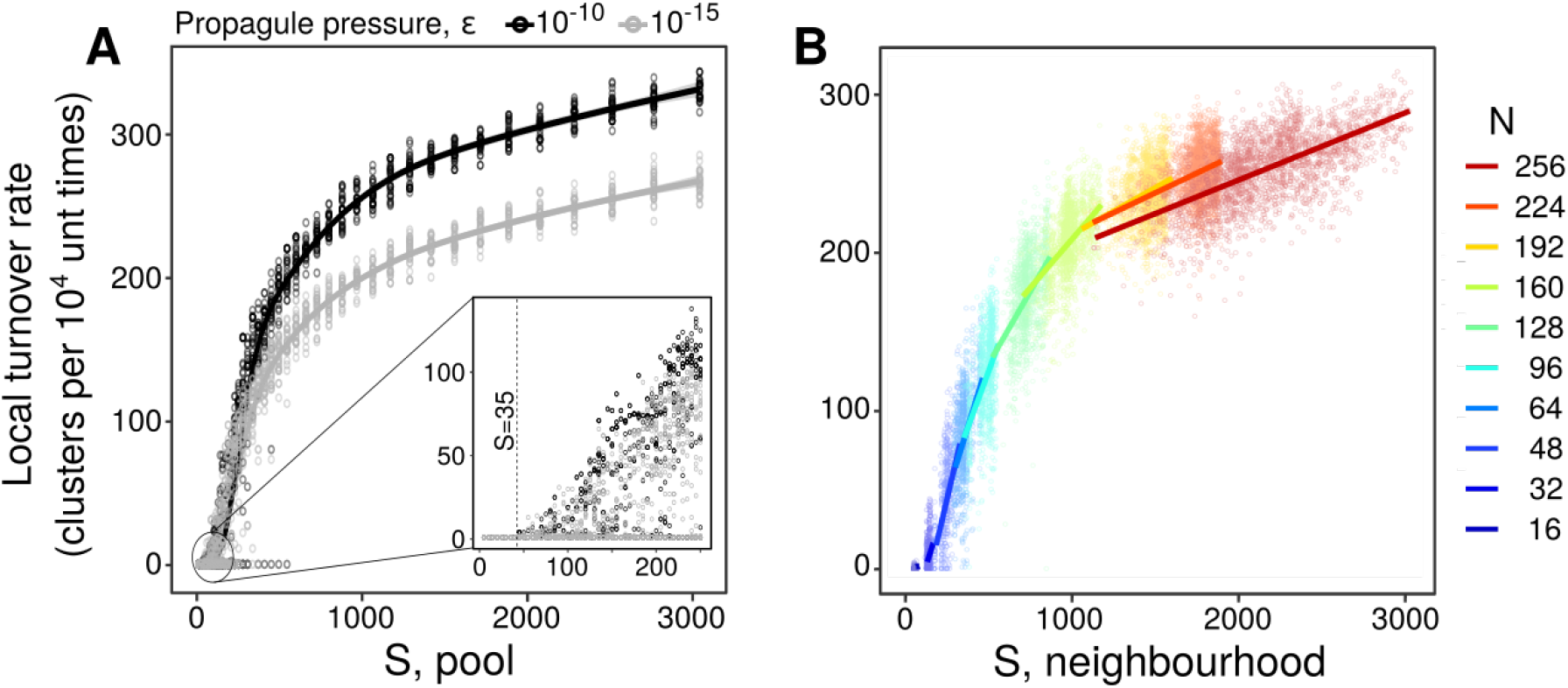
Ecological mass effects drive autonomous turnover. **A**: The number of compositional clusters detected, plotted against the size of the pool of potential invaders for an isolated LV community using a propagule pressure *ϵ* of 10^−10^ and 10^−15^, fit with a generalized additive model^53^. For *S* < 35 a single cluster is detected. For *S* ≥ 35 autonomous turnover occurs (≥ 1 compositional clusters) with the transition indicated by the dashed line (inset). **B**: Qualitatively identical behaviour was observed for model metacommunities in which ‘propagule pressure’ arises due to ecological mass effects from the local neighbourhood. Each point represents a single patch. Lines in **B** are standard linear regressions. The good alignment of subsequent fits demonstrates that neighbourhood diversity is the dominating predictor of cluster number, rather than patch number *N. A_ij_* = 0.5 with probability 0.5, *φ* = 10, *σ*^2^ = 0.01, *ℓ* = 0.5.

There is thus a close correspondence between dynamically isolated LV models and LVMCM metacommunities in the sequence of dynamic states as propagule richness increases and in the resulting complexity of dynamics quantified by counting compositional clusters. This suggests that underlying heteroclinic networks, which are revealed by adding propagule pressure in isolated communities, explain the complex dynamics seen in LVMCM metacommunities.

For the isolated LV community, the threshold beyond which autonomous turnover is detected (> 1 compositional cluster) occurs at a pools size of around *S* =35 species, consistent with the theoretical prediction^50^ of the transition between the UFP and MA phases (supplementary text). Close inspection of this threshold reveals an important and hitherto unreported relationship between the transition into the MA phase and local ecological limits set by the onset of ecological structural instability, which is known to regulate species richness in LV systems subject to external invasion pressure^23,24^: in Supplementary Material we show that the boundary between the UFP and MA phases^50^ coincides precisely with the onset of structural instability^24^ (Eqs. S6-S12). For LVMCM metacommunities, the relationship revealed analytically in the Supplementary Material is numerically confirmed in Fig. 5. During assembly, local species richness increases until it reaches the limit imposed by local structural instability. Further assembly occurs via the ‘regionalisation’ of the biota^54^–a collapse in average range sizes^23^ and associated increase in spatial beta diversity-until regional diversity limits are reached^23^. The emergence of autonomous turnover coincides with the onset of species saturation *at the local scale*. Autonomous turnover can therefore serve as an indirect indication of intrinsic biodiversity regulation via local structural instability in complex communities.

**Figure 5.**
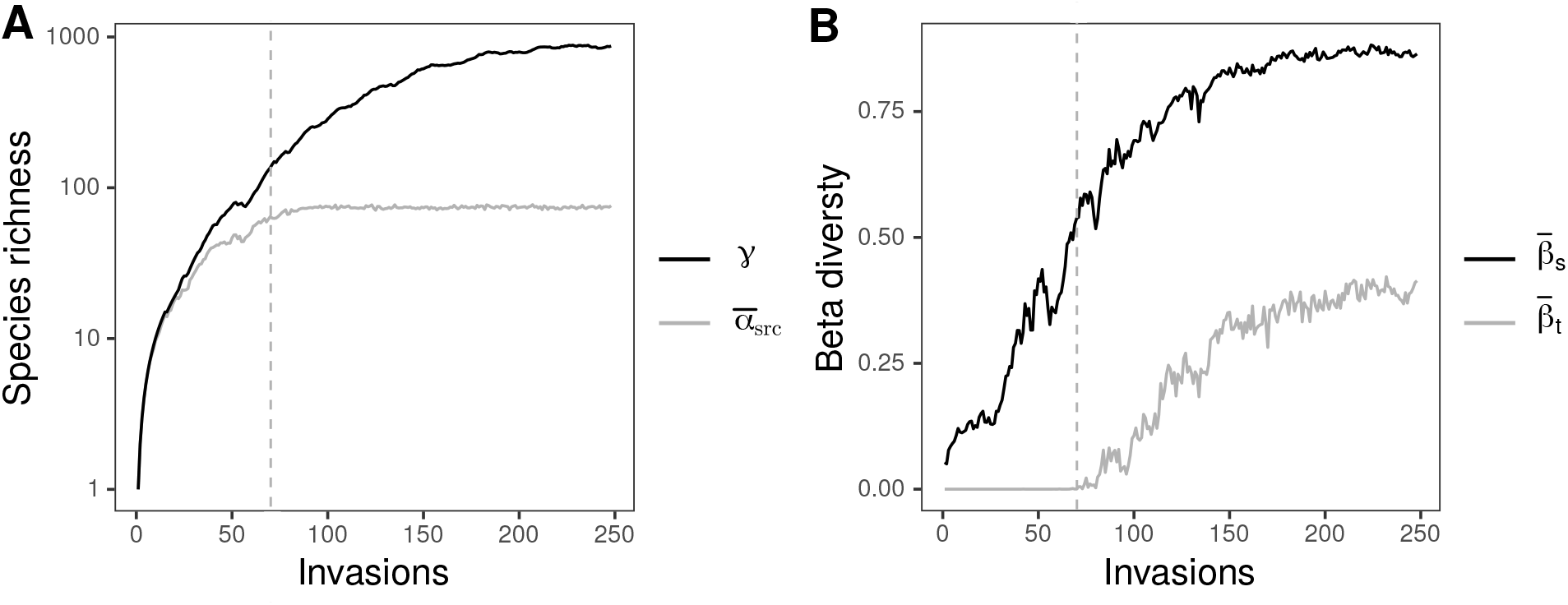
The emergence of temporal turnover during metacommunity assembly. **A**: local species richness, defined by reference to source populations only (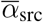, grey) and regional diversity (black) for a single metacommunity of *N* = 32 coupled communities during iterative invasion of random species. We quantify local source diversity 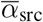 as the metacommunity average of the number *α*_src_ of non-zero equilibrium populations persisting when immigration is switched off (off-diagonal elements of **D**set to zero), since this is the component of a local community subject to strict ecological limits to biodiversity. Note the log scale chosen for easy comparison of local species richness and regional diversity. **B**: Increases in regional diversity beyond local limits arise via corresponding increases in spatial turnover (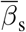, black). Autonomous temporal turnover (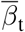, grey) sets in precisely when average local species richness 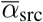 has reached its limit, reflecting the equivalence of the transition to the MA phase space and the onset of local structural instability. In both panels, the dashed line marks the point at which autonomous temporal turnover was first detected. *A_ij_* = 0.3 with probability 0.3, *φ* = 10, *σ*^2^ = 0.01, *ℓ* = 0.5. Both spatial and temporal turnover computed as the mean BC dissimilarity.

Thus, we have shown that propagule pressure perturbs local communities away from unstable equilibria and drives compositional change. In order to invade, however, species need to be capable of passing through biotic and abiotic filters^33–35^. We would expect, therefore, that turnover would be suppressed in highly heterogeneous or poorly connected environments where mass effects are weak. Indeed, by manipulating the autocorrelation length *φ*, and variance *σ*^2^ of the abiotic filter represented by the matrix **R** and the characteristic dispersal length *ℓ*, we observe a sharp drop-off in temporal turnover in parameter regimes that maximise between-patch community dissimilarity (short environmental correlation or dispersal lengths Fig. S10).

### The macroecology of autonomous turnover

We find important similarities between temporal and spatio-temporal biodiversity patterns emerging in model metacommunities in the absence of external abiotic change and in empirical data (Fig. 6), with quantitative characteristics lying within the ranges observed in natural ecosystems.

**Figure 6.**
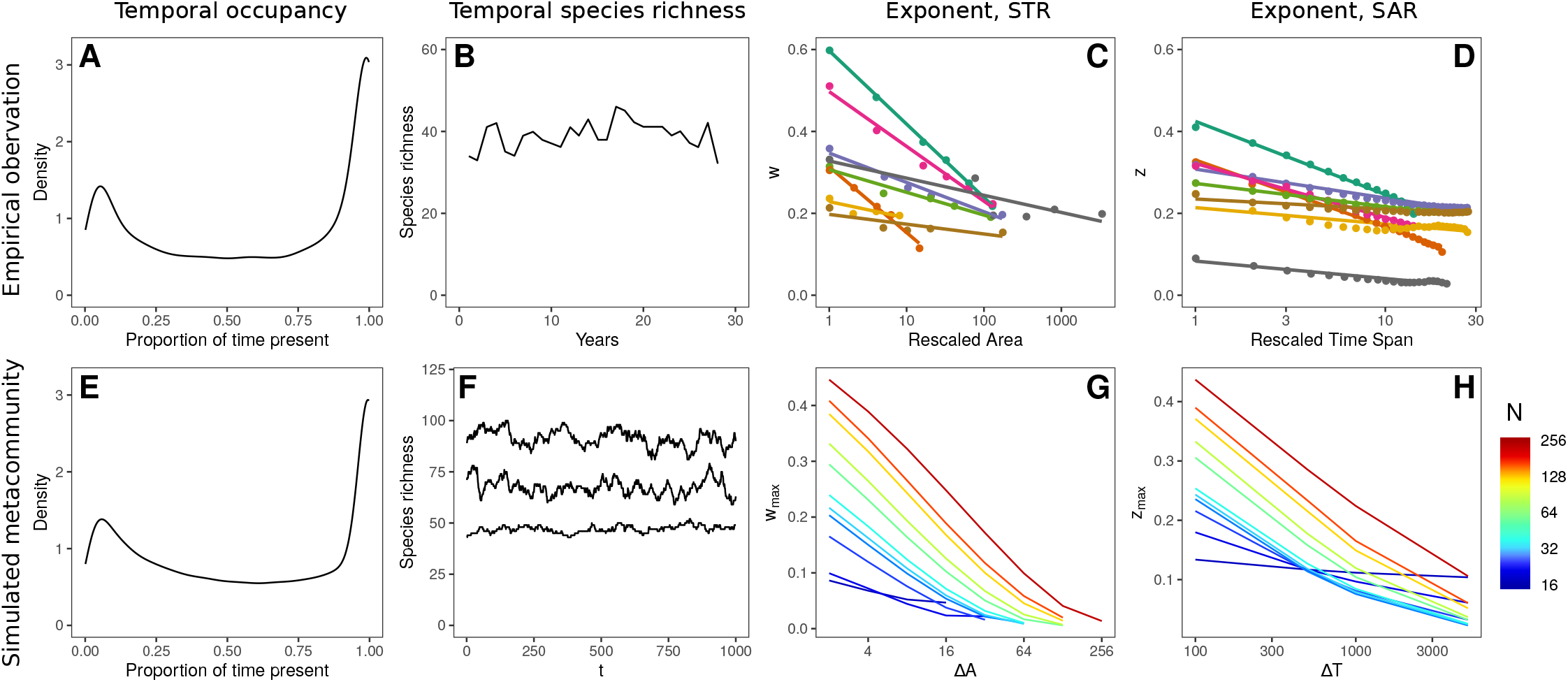
Macroecological signatures of autonomous compositional change. A bimodal distribution in temporal occupancy observed in North American birds^55^ (**A**) and in simulations (**E**, *N* = 64, *φ* = 5, *σ*^2^ = 0.01, *ℓ* = 0.5). Intrisically regulated species richness observed in estuarine fish species^60^ (**B**) and in simulations (**F**, *N* = 64, *φ* = 5, *σ*^2^ = 0.01, *ℓ* = 0.5). The decreasing slopes of the STR with increasing sample area^14^ (**C**), and the SAR with increasing sample duration^14^ (**D**) for various communities and in simulations (**G**and **H**, *N* = 256, *φ* = 10, *σ*^2^ = 0.01, *ℓ* = 0.5). In **C** and **D** we have rescaled the sample area/duration by the smallest/shortest reported value and coloured by community (see original study for details). In **G** and **H** we study the STAR in metacommunities of various size *N*, represented by colour. Limited spatio-temporal turnover in the smallest metacommunties (blue colours) greatly reduces the exponents of the STAR relative to large metacommunities (red colours). *A_ij_* = 0.5 with probability 0.5 in all cases.

#### Temporal occupancy

The proportion of time in which species occupy a community tends to have a bi-modal empirical distribution^55–57^ (Fig. 6A). The distribution we found in simulations (Fig. 6E) closely matches the empirical pattern.

#### Community structure

Temporal turnover has been posited to play a stabilizing role in the maintenance of community structure^58,59^. In an estuarine fish community^60^, for example, species richness (Fig. 6B) and the distribution of abundances were remarkably robust despite changes in population biomasses by multiple orders of magnitude. In model metacommunities with autonomous turnover we found, likewise, that local species richness exhibited only small fluctuations around the steady-state mean (Fig. 6F, three random local communities shown) and that the macroscopic structure of the community was largely time invariant (Fig. S11). In the light of our results, we propose the absence of temporal change in community properties such as richness or the abundance distribution despite potentially large fluctuations in population abundances^60^ as an indication of predominantly autonomous compositional turnover.

#### The Species-Time-Area-Relation, STAR

The species-time-relation (STR), typically fit by a power law of the form *S ∝ T^w^*^14,61,62^, describes how observed species richness increases with observation time *T*. The exponent *w* of the STR has been found to be remarkably consistent across taxonomic groups and ecosystems^14,15,63^, indicative of some general population dynamical mechanism. However, the exponent of the STR decreases with increasing sampling area^14^, and the exponent of the empirical Species Area Relation (SAR) (*S ∝ A^z^*) consistently decreases with increasing sampling duration^14^ (Fig. 6C, D). We tested for these patterns in a large simulated metacommunity with *N* = 256 patches by computing the STAR for nested subdomains and variable temporal sampling windows (see Methods). We observed exponents of the nested SAR in the range *z* = 0.02-0.44 and for the STR a range *w* = 0.01-0.44 (Fig. S12), both in good agreement with observed values^15,64^. We also found a clear decrease in the rate of species accumulation in time as a function of sample area and vice-versa (Fig. 6G, H).

Thus, the distribution of temporal occupancy, the time invariance of key marcoecological structures and the STAR in our model metacommunities match observed patterns. This evidence suggests that such autonomous dynamics cannot be ruled out as an important driver of temporal compositional change in natural ecosystems.

## CONCLUSIONS

Current understanding of the mechanisms driving temporal turnover in ecological communities is predominantly built upon phenomenological studies of observed patterns^2,65–67^ and is unquestionably incomplete^8,60^. That temporal turnover can be driven by external forces—e.g. seasonal or long term climate change, direct anthropogenic pressures—is indisputable. A vitally important question is, however, how much empirically observed compositional change is actually due to such forcing. Recent landmark analyses of temporal patterns in biodiversity have detected no systematic change in species richness or structure in natural communities, despite rates of compositional turnover greater than predicted by stochastic null models^1,68–70^. Here we have shown that empirically realistic turnover in model metacommunities can occur via precisely the same mechanism as that responsible for regulating species richness at the local scale. While the processes regulating diversity in natural communities remain poorly understood, our theoretical work suggests local structural instability may explain these empirical observations in a unified and parsimonious way. Therefore, we advocate for the application of null models of metacommunities dynamics that account for natural turnover in ecological status assessments and predictions based on ancestral baselines.

How do the turnover rates that we find in our model compare with those observed? Our current analytic understanding of autonomous turnover is insufficient for estimating the rates directly from parameters, but the simulation results provide some indication of the expected order of magnitude, that can be compared with observations. Key for such a comparison is the fact that, because the elements of **R** are 1 on average, the time required for an isolated single population to reach carrying capacity is 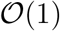 unit times. Fig. S11B suggests that transitions between community states occur at the scale of around 10-50 unit times. This gives a holistic, rule-of-thumb estimate for the expected rate of autonomous turnover, depending on the typical reproductive rates of the guild of interest. In the case of macroinvertebrates, for example, the time required for populations to saturate in population biomass could be of the order of a month or less. By our rule of thumb, this would mean that autonomous community turnover would occur on a timescale of years. In contrast, for slow growing species like trees, where monoculture stands can take decades to reach maximum population biomass, the predicted timescale for autonomous turnover would be on the order of centuries or more. Indeed, macroinvertebrate communities have been observed switching between community configurations with a period of a few years^71,72^, while the proportional abundance of tree pollen and tree fern spores fluctuates in rain forest bog deposits with a period of the order of 10^3^ years^73^—suggesting that predicted turnover rates are biologically plausible.

Our simulations revealed a qualitative transition from ‘small’ metacommunities, where autonomous turnover is absent or minimal, to ‘large’ metacommunities with pronounced autonomous turnover (Fig. 2). The precise location of the transition between these cases depends on details such as dispersal traits, the ecological interaction network, and environmental gradients (Fig. S3). Taking, for simplicity, regional species richness as a measure of metacommunity size suggests that both ‘small’ and ‘large’ communities in this sense are realised in nature. In our simulations, the smallest metacommunities sustain 10s of species, while the largest have a regional diversity of the order 10^3^, which is not large comparable to the number of tree species in just 0.25 km^2^ of tropical rainforest (1,100 – 1, 200 in Borneo and Ecuador^74^) or of macroinvertebrates in the UK (> 32,000 ^75^). Within the ‘small’ category, where autonomous turnover is absent, we would therefore expect to be, e.g. communities of marine mammals or large fish, where just a few species interact over ranges that can extend across entire climatic niches, implying that the effective number of independent “patches” is small and providing few opportunities for colonisation by species from neighbouring communities. Likely to belong to the ‘large’ category are communities of organisms that occur in high diversity with range sizes that are small compared to climatic niches, such as macroinvertebrates. For these, autonomous turnover of local communities can plausibly be expected based on our findings. Empirically distinguishing between these two cases for different guilds will be an important task for the future.

For metacommunities of intermediate spatial extent, autonomous turnover is characterized by sharp transitions between cohesive states at the local scale. To date, few empirical analyses have reported such coherence in temporal turnover, perhaps because the taxonomic and temporal resolution required to detect such patterns is not yet widely available. Developments in biomonitoring technologies^76^ are likely to reveal a variety of previously undetected ecological dynamics, however and by combining high resolution temporal sampling and metagenetic analysis of community composition, a recent study demonstrated cohesive but short-lived community cohorts in coastal plankton^77^. Such Clementsian temporal turnover may offer a useful signal of autonomous compositional change in real systems.

Thus, overcoming previous computational limits to the study of complex metacommunities^10,78^, we have discovered the existence of two distinct phases of metacommunity ecology—one characterized by weak or absent autonomous turnover, the other by continuous compositional change even in the absence of external drivers. By synthesizing a wide range of established ecological theory^10,23,24,46,50,51^, we have heuristically explained these phases. Our explanation implies that autonomous turnover requires little more than a diverse neighbourhood of potential invaders, a weak immigration pressure, and a complex network of interactions between co-existing species.

## Acknowledgements

We thank Lars Chittka, Laurent Frantz and three anonymous referees for comments on earlier drafts of this paper.

## Funding

This work forms part of the project “Mechanisms and prediction of large-scale ecological responses to environmental change” funded by the Natural Environment Research Council (NE/T003510/1).

## Author contributions

AGR conceived of the study. JDO and AGR designed the model. JDO developed the model, performed simulations, analysed the data and drafted the manuscript. All authors interpreted model outputs in comparison with observations and contributed to manuscript writing.

## Competing interests

The authors declare no competing interests.

## Data and materials availability

Should this manuscript be accepted simulation data supporting the results will be archived in a public repository and the data DOI will be included at the end of the article.

## SUPPLEMENTARY MATERIALS

Materials and Methods

Supplementary text

Figs. S1 – S10

## MATERIALS AND METHODS

### Metacommunity assembly

The dynamics of local population biomasses *B_ix_*(*t*) were modelled using a spatial extension to the multispecies Lotka-Volterra competition model^23^:

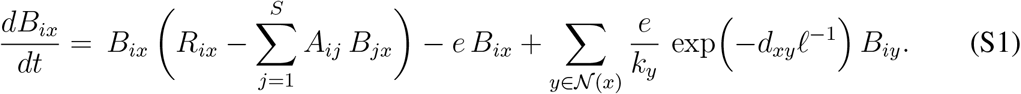

The competitive coupling coefficients *A_ij_* for *i ≠ j* were sampled from discrete distributions. Generally, *A_ij_* were set to 0.5 with a probability of 0.5 and to 0 otherwise, however, for the simulation shown in Fig. 5, we relaxed the dynamic coupling and instead set *A_ij_* to 0.3 with a probability of 0.3. This delayed the onset of local structural instability during metacommunity assembly, making the coincident emergence of local biodiversity regulation and autonomous compositional turnover visually clearer.

Environmental heterogeneity was modelled implicitly through spatial variation in species’ intrinsic growth rates *R_ix_*. Specifically, the *R_ix_* were sampled independently for each species *i* from a Gaussian random field^85^ with mean *μ* = 1.0 and standard deviation *σ*, generated via spectral decomposition^86^ of the *N* × *N* landscape covariance matrix with elements ∑_*xy*_ = exp [−*φ*^−1^*d_xy_*], where *d_xy_* denotes the Euclidean distances between patches *x* and *y*, and *φ* the autocorrelation length (Fig. S2).

The dispersal matrix **D** (Eq. (1)) has diagonal elements *D_xx_* of −*e*, where *e*, the fraction of biomass leaving patch *x* per unit time, was kept fixed at 0.01 for all simulations. For pairs of patches connected by an edge in the spatial network, the immigration terms were modelled as negative exponentials 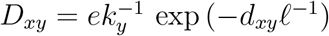, controlled by a dispersal length parameter *ℓ*, thus assuming a propensity for propagules to transition to nearby sites. The normalisation constant *k_y_* divides the biomass departing patches *y* between all other patches in *its* local neighbourhood 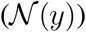, weighted by the ease of reaching each patch i.e. 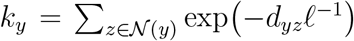, implying an active dispersal process.

Metacommunities were assembled through a stepwise invasion process (Fig S1). In each iteration of the algorithm, 0.05S + 1 new species were introduced to the metacommunity, with *S* denoting the current extant species richness. The invaders were tested to ensure positive growth rates at low abundance. This was done by introducing a multiple of 0.05S + 1 newly generated species into all patches at very low abundance, then simulating for a handful of time steps and testing for increasing biomass trajectories in at least one patch. Of the successful invaders, 0.05S + 1 were randomly selected and each introduced at 10^−6^ biomass units into the patch in which its growth rate was greatest during testing. After invaders were introduced, metacommunity dynamics were simulated using the SUNDIALS^80^ numerical ODE solver. The time between invasions we kept fixed at 500 unit times, and before each new invasion the metacommunity was scanned and species with biomass smaller than 10^−4^ biomass units in all patches of the network were considered regionally extinct and removed from the model. The assembly algorithm aims to remove all species whose total biomass declines to zero in the course of the system’s complex dynamics. In rare cases autonomous fluctuations may drive one of the remaining species to very low abundance in all patches, however the majority retain local biomass above the detection threshold in at least one patch at all times.

To assemble models of sufficient spatial extent and species richness, we developed a parallel implementation of the assembly model that makes use of the algorithmic domain decomposition method^81^ for the population-dynamical simulations. This involves decomposing the metacommunity into spatial subdomains of equal numbers of patches, each of which is simulated by a unique parallel process (CPU), with boundary states regularly broadcast between processes. The code was run on the Apocrita high-performance cluster at Queen Mary, University of London^87^. This permitted assembly of saturated metacommunities of up to *N* = 256 patches harbouring *S* ~ 3000 species, thus breaking through frequently lamented computational limits^10,78^ on the numerical study of metacommunities.

### Quantifying autonomous turnover

For fully assembled metacommunities, we simulated and stored time series of *t*_max_ = 10^4^ metacommunity samples *B_ixt_* = *B_ix_*(*t*) taken in intervals of one unit time. In these metacommunity timeseries, we measured spatio-temporal turnover based on i) compositional dissimilarity, ii) the distribution of temporal occupancy, iii) the number of compositional clusters detected using hierarchical clustering, and iv) via species accumulation curves generated using sliding spatial and temporal sampling windows. Metrics were selected in order to answer specific questions, or for comparison to observed patterns. Some analyses require quantifying local species richness. This was done by setting a detection threshold of 10^−4^ biomass units, below which populations are considered absent from the community. *Local source diversity*, which we define in Fig. 5, is a related but different diversity measure that is more adequate for quantifying the component of a local community subject to local ecological limits to biodiversity.

### Compositional dissimilarity

Spatial/temporal compositional similarity was quantified using the Bray-Curtis^42^ similarity index via the function `vegdist` in the R package “vegan”^91^.

### Temporal occupancy

We assessed temporal occupancy by first converting biomass into presence-absence data (*P_ixt_* = 1 for all *B_ixt_* > 10^−4^, and 0 otherwise). Then, for all populations present at least once, we computed the temporal occupancy (*TO_ix_*) as the proportion of the time interval of length *t*_max_ during which that population was present:

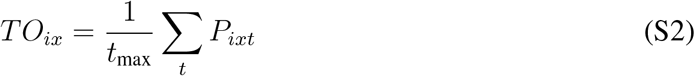

### Hierarchical clustering

We assessed the degree of temporal clustering in community composition using complete linkage hierarchical clustering^92^ of the Bray-Curtis similarity matrix, which gives an approximate measure of the number of unstable equilibria between which the dynamical system fluctuates. We computed the number of clusters using a threshold of 75% similarity, which reflects the structure visible in pairwise dissimilarity matrices (Fig. S5A and B).

### Spatio-temporal species accumulation

We studied the STR and SAR in model metacommunities using a sliding window approach, asking, for given 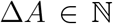 and 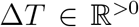, how many species *S*^obs^ were detected on average in sets 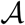 of 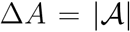 patches during any time interval 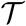 of Δ*T* unit times length. Specifically, for a metacommunity of *N* = 2^8^ = 256, the spatial windows were Δ*A* ∈ {2^0^, 2^1^,…, 2^8^} patches, while the temporal windows were Δ*T* ∈ {1, 5, 10, 50, 100, 500, 1000} unit times. For each patch *x* ∈ {1,…, *N*} the spatial subsample was then defined as the set 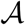 consisting of the focal patch and its Δ*A* − 1 nearest neighbours. Similarly, for each *t* ∈ {1,…, *t*_max_ − Δ*T*} the sliding temporal window 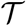 was defined as the ΔT successive recording time steps in the range *t* to *t* +Δ*T*. The species richness observed in a given spatio-temporal sub-sample was then computed as

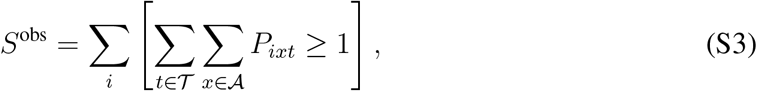

where the Iverson brackets [.] denote the indicator function ensuring species are counted only once. Finally, the average of *S*^obs^ for a given spatio-temporal sample size was computed in all combinations.

In closed systems, the species accumulation in both space and time must ultimately saturate, either when the entire metacommunity or entire time series is sampled. Thus we defined the exponents *z* and *w* of the STAR as the maximum slopes of the SAR/STR on double logarithmic axes (Fig. S12).

## SUPPLEMENTARY TEXT

### Spatial parameterization

Other than patch number *N*, the parameters that most impact the spatio-temporal structure of model metacommunities are the environmental correlation length *φ*, the variability of the environment *σ*^2^, and the dispersal length *ℓ*. In order to understand the role of these parameters for autonomous turnover, we fixed *N* = 64 and assembled metacommunity models with *σ*^2^, ℓ ∈ {1 × 10^−2^, 5 × 10^−2^, 1 × 10^−1^, 5 × 10^−1^, 1}, and φ∈{1, 5, 10, 50, 100}in all combinations and computed the resulting temporal beta diversity as the mean spatially averaged temporal BC dissimilarity observed in 10 replicates of each parameterization. Rates of autonomous turnover varied in a complex but systematic way under variation in the spatial parameterization of the model, with turnover being weakly correlated with the dispersal length and maximized for intermediate habitat heterogeneity and autocorrelation (Fig. S3). Weak abiotic heterogeneity seeds the non-uniform spatial structure of the metacommunity and therefore promotes turnover. For large enough spatial networks, dispersal limitation and competitive repulsion alone are sufficient to drive autonomous dynamics in perfectly uniform landscapes. The scan of the parameter space allowed selection a parameterization with strong autonomous turnover: *φ* = 10, *σ*^2^ = 0.01, *ℓ* = 0.5 (peak in Fig. S3A). Using this combination of parameters we then assembled metacommunity models of *N* = 8, 16, 32, 48, 64, 80, 96, 128, 160, 192, 224, 256 patches.

To some extent, the complex roles of parameters *φ, σ*^2^, and *ℓ*, shown in Fig. S3, can be distilled into the effect on a single parameter: the average spatial community dissimilarity at the local neighbourhood scale. This is due to the fact that the impact of each of the parameters, which control the between-patch differences in environment and the strength of mass effects, is reflected in the degree of spatial beta diversity within the metacommunity. To demonstrate this we used the multiple-site dissimilarity metric derived in Ref.^83^, which generates an unbiased total beta diversity metric for systems of three or more sites/time points. Since both local neighbourhood and (correspondingly) temporal turnover vary within a given metacommunity, we show the beta diversity metrics averaged over all patches.

Temporal turnover responded unimodally to local neighbourhood dissimilarity (Fig. S10) over the parameter range of Fig. S3, suggesting that spatial parameterizations that maximise *β_s_*, either through exaggerating abiotic differences between adjacent local communities or dampening mass effects, can *elevate* neighbourhood diversity while simultaneously *suppressing* the pool of species that can actually invade.

This result makes plausible why empirical studies have detected a range of statistical associations between spatial and temporal turnover in natural ecosystems. Positive, negative, unimodal, and non-significant relationships have been reported between temporal turnover and species richness or spatial turnover^15,95–99^. The unimodal response shown in Fig. S10 may help to resolve these apparent contradictions: it is not species richness or spatial dissimilarity *per se* that best predict temporal turnover, but the size of the pool of species capable of passing through biotic and abiotic filters to invade a local community.

### Phase space of a generalised Lotka-Volterra community

Analytic theory^50^ predicts a sharp transition between what has been called the Unique Fixed Point (UFP) and Multiple Attractor (MA) phases. In Fig. S6 we reproduce the phase portrait for such a system and note that our explicitly modelled metacommunities reveal a gradual transition in the MA phase space from oscillatory, to Clementsian and into Gleasonian turnover regimes. Assuming large *S*, the sharp transition between UFP and MA phases has been shown^50^ to occur at species richness

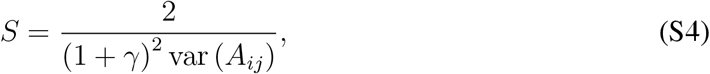

where *γ* = corr (*A_ij_, A_ji_*) denotes the degree of correlation in the effects two species have on each other, measuring the symmetry of interspecific interaction strengths, and var (*A_ij_*) is the variance in the distribution. In our model we use a random interaction matrix for which *γ* = 0. We sample interaction coefficients from a discrete distribution with var (*A_ij_*) = (0.25)^2^ giving a predicted transition into the MA phase space at *S* = 32 species. Thus, while the prediction is approximate for small *S* communities with non-uniform intrinsic growth rates, a numerically observed threshold of around 35 species in the isolated LV model (Fig. 4C inset) is consistent with these analytic predictions.

### Isolated LV communities

To explore the emergence of heteroclinic networks in LV models, we studied an isolated LV model with and without coupling to an implicitly modelled neighbourhood species pool. The dynamics of the model follow

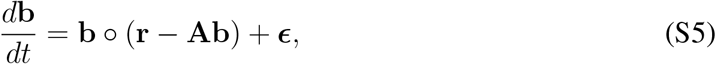

where **b** is a population biomass vector of length *S*, **r** is a vector of independent random normal variables with mean 1 and variance *σ*^2^ = 0.01 representing maximum intrinsic growth rates, **A** is a competitive overlap matrix and the vector **e** represents the slow immigration of biomass corresponding to a weak propagule pressure. The elements *ϵ_i_* are analogous the to explicitly modelled immigration terms *B_ix_D_xy_* of the full metacommunity model.

As in the metacommunity model, interspecific competition coefficients *A_ij_* were set to 0.5 with a probability of 0.5 for *i ≠ j* and otherwise to zero, while *A_ii_* = 1, for all *i*. We enforced *b_i_* > 0 for all *i* by simulating dynamics in terms of logarithmic biomass variables. In simulating this model, we did not follow the common practice of removing species whose biomass drops below some threshold. Instead all species were retained. We consider two situations: with and without the inclusion of a weak propagule pressure **ϵ**.

### Heteroclinic networks in the case without propagule pressure

We first demonstrate in simulations that, indeed, as predicted under certain constraints^46^, stable heteroclinic networks exist in the MA phase of model Eq. (S4) for ***ϵ*** = 0. For this we choose *S* = 300, which, with other parameters set as described above, brings us deeply into the MA phase of the model. Simulations were initialised by setting all *B_i_* = 10^−3^ (1 ≤ *i ≤ S*) at *t* = 0. The system was simulated until *t* = 2.1·10^7^ and system states recorded at times *t* = 2.1 · 10^*j*/1000^(0 ≤ *j* ≤ 7000). As illustrated in Fig. S7, while dynamics tend to become slower for larger *t*, no stable equilibrium or other simple attractor appears to be ever reached—as expected for a system approaching a heteroclinic network. Instead, as expected when a heteroclinic network exists, the system bounces around between unstable equilibria, apparently in a random fashion. Unexpected to us, however, the system appears to visit not only unstable equilibria in its transient, but occasionally also unstable periodic orbits (*t* ≈ 1.3 · 10^4^ in Fig. S7) and perhaps more complex invariant sets (*t* ≈ 1.2 · 10^6^ in Fig. S7).

One might wonder whether there is any tendency for dynamics to eventually come to a halt. To study this question, we calculated the number of changes in community composition (species colonisations and extinctions) between all pairs of subsequently recorded system states, where we considered a species *i* as “present” if *B_i_* > 10^−4^, and from this the momentary rate of change in composition on the ln(t) scale by dividing by ln(10^1/1000^). In Fig. S8 we show the time series of the centred moving average over this number for 100 subsequent pairs or recordings, and averages for non-overlapping adjacent blocks for 300 pairs. Spikes where the rate of change is particularly high correspond to brief phases of regular or irregular oscillation. We performed a median regression of the block-wise averages by a power law of the form: (rate) ~ *t^ν^*. Median regression was used to de-emphasize the spikes. For the simulation shown in Fig. S7 we found that *v* did not differ significantly from zero, implying a decline of the turnover rate on the natural time axis as *t*^−1^. When we repeated this analysis for 15 independent simulations (two of which failed due to numerical issues), we observed a tendency for *ν* to be slightly positive (*ν* = 0.054 ± 0.020, t-test *t* = 2.67, *p* = 0.020), perhaps because the effect of oscillatory phases on the mean turnover rate on the ln(*t*)-scale increases with increasing t. Overall, however, the decline of turnover rate approximately as *t*^−1^ was confirmed, providing evidence for the existence of an attracting heteroclinic network that the LV system Eq. (S5) with ***ϵ*** = 0 slowly approaches.

Use of logarithmic biomass variables was essential for these simulations. We found that median species biomass at the end of each run was typically around 10^−3,500,000^, much smaller than the smallest number representable by double precision floating point arithmetic, which is around 2 · 10^−308^. Needless to say, these small numbers mean that the simulations with *ϵ* = 0 are, while instructive, ecologically unrealistic.

### Heteroclinic networks in the case with propagule pressure

The case ***ϵ*** > 0, where dynamics move alongside the underlying heteroclinic network without ever fully approaching it, is discussed in the Main Text as it provides a useful intermediate between the explicit metacommunity model and the more tractable isolated community. In Fig. S9 we show that the transition from oscillatory to Clementsian and finally Gleasonian turnover regimes can also be observed in these isolated LV models (*ϵ_i_* = *ϵ* = 10^−15^ for all *i*, other parameters as above).

### Local structural instability drives autonomous turnover

Species richness in competitive LV communities is intrinsically limited by the onset of ecological structural instability. Here we show analytically that for isolated communities the boundary between the UFP and MA phases^50^ is identical to the structurally unstable limit^24^.

The transition between UFP and MA phase for competitive LV models occurs^50^ when

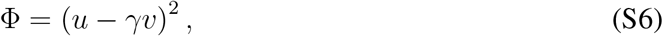

where Φ:= *S*/S* is the proportion of species persisting, i.e. the ratio between the number *S*\* of species that persist and the pool size *S*, and again *γ* = cor (*A_ij_, A_ji_*). The quantities *u* and *v* in Eq. (S6) are given by

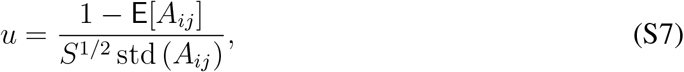

with E[*A_ij_*] and std (*A_ij_*) denoting mean and standard deviation of the distribution of off-diagonal entries of **A**, respectively, and

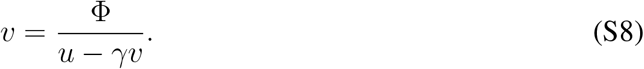

For *γ* ≠ 0, Eq. (S8) does not have a unique solution for *v*. The equivalent quadratic equation *γv*^2^ − *uv* + Φ = 0 has two solutions, one of which diverges as *γ* → 0; this we discard. The other solution is

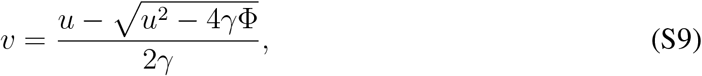

which becomes *v* = Φ/*u* for *γ* → 0, consistent with Eq. (S8). Substitution of Eq. (S9) into
Eq. (S6) gives

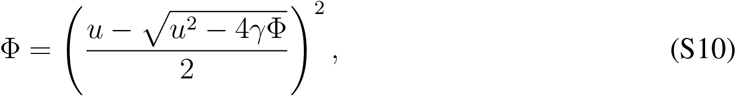

which can be shown in a standard calculation to be equivalent to

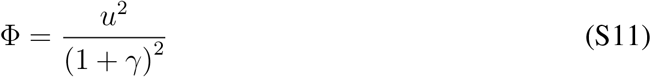

for *u* > 0 and −1 < *γ* < 1. Finally, substituting Eq. (S7) into Eq. (S11) gives

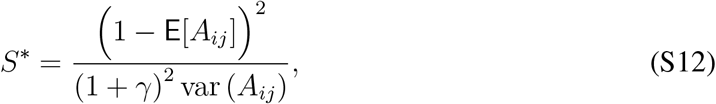

which is exactly the theoretical limit of structural instability in isolated LV communities [Eq. (18.3) of Ref. 24], thus demonstrating that UFP-MA phase boundary and the onset of structural instability perfectly coincide.

### Temporal patterns in community structure

Fluctuations in local population biomasses as communities move between unstable equilibria in heteroclinic networks can span multiple orders of magnitude (red trajectories in Fig. S11A) and lead to significant temporal turnover in community composition (Fig. S11B). In contrast, the high-level properties of the assemblages remain largely unchanged. This is evident in the dampening of biomass fluctuations at metapopulation and metacommunity scales via a spatial portfolio effect^58,59,82^ (blue and black trajectories in Fig. S11A), but also in the robustness of species biomass distribution (Fig. S11C) and range size distribution (Fig. S11D, range sizes computed as in Ref.^23^). In this case the mean relative biomass and range size are plotted irrespective of species identity (black lines) along with the mean ±one standard deviation (grey lines), for direct comparison with Ref.^60^. The relatively small standard deviations demonstrate a temporally robust distribution of metapopulation biomasses and spatial ranges, despite large fluctuations at the local scale.

### STAR in large metacommunity models

We characterised the within assemblage STAR using a moving spatio-temporal window as described in the main text and comparing the resulting SAR and STR exponents. In Fig. S12 we show the nested SAR and STR for a single metacommunity of *N* = 256. The number of species detected for large spatial or temporal windows necessarily saturates in closed systems. We therefore defined the exponents of the STAR, displayed in Fig. 6 of the main text, as the maximum slope of the SAR/STR on double logarithmic axes.

## Supplementary figures

**Figure S1.**
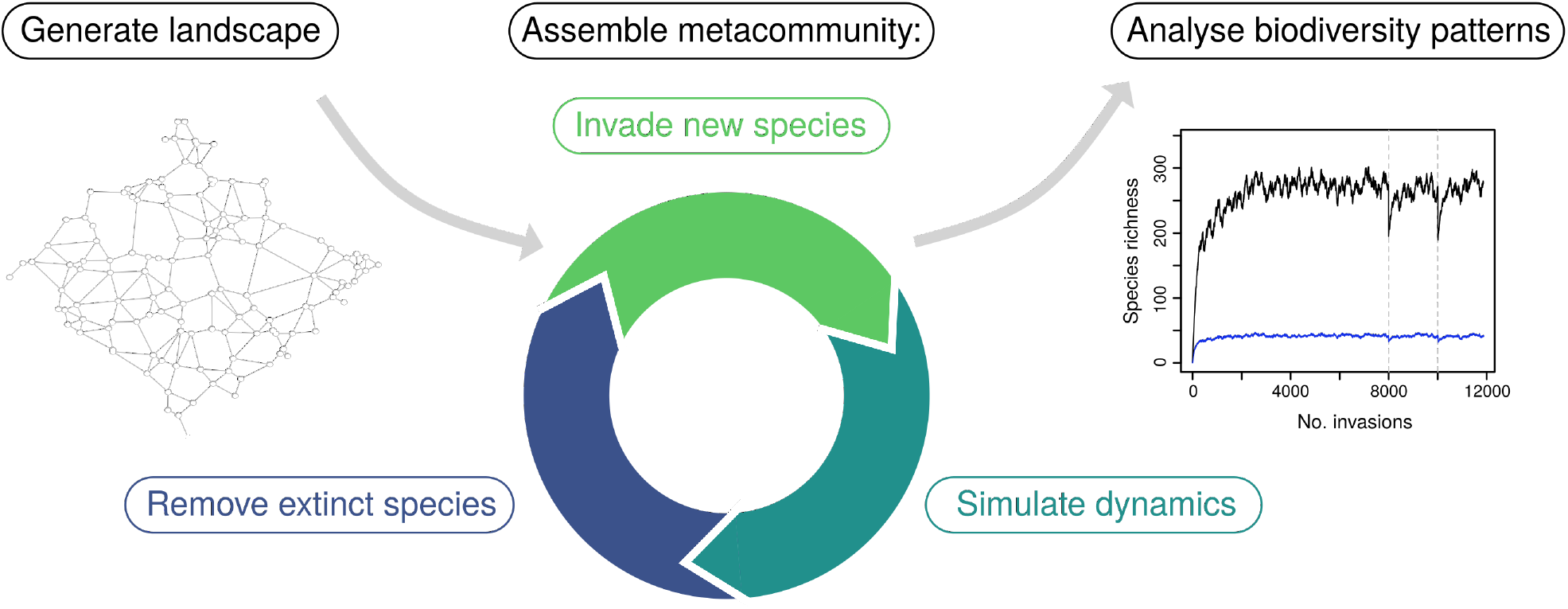
The metacommunity assembly algorithm. First, a random planar graph is generated with spatial coordinates sampled at random and patches connected via the Gabriel^40^ algorithm. Communities are then assembled iteratively: species are generated with intrinsic growth rates and interaction coefficients sampled from random distributions, introduced into the metacommunity at low abundance, metacommunity population-dynamics are simulated, and regionally extinct species are removed from the model before the next iteration. Eventually the metacommunity reaches both its local and regional diversity limits, the situation studied in the main text. In the inset a single metacommunity assembly process is shown; the black line represents regional species richness, the blue line average local species richness. Both of which are intrinsically regulated, as demonstrated by the effect of random removals of species (dashed lines) and subsequent re-assembly: local richness is barely affected and regional richness returns to the approximate same level. Inset adapted from Ref.^23^. See text for detailed description.

**Figure S2.**
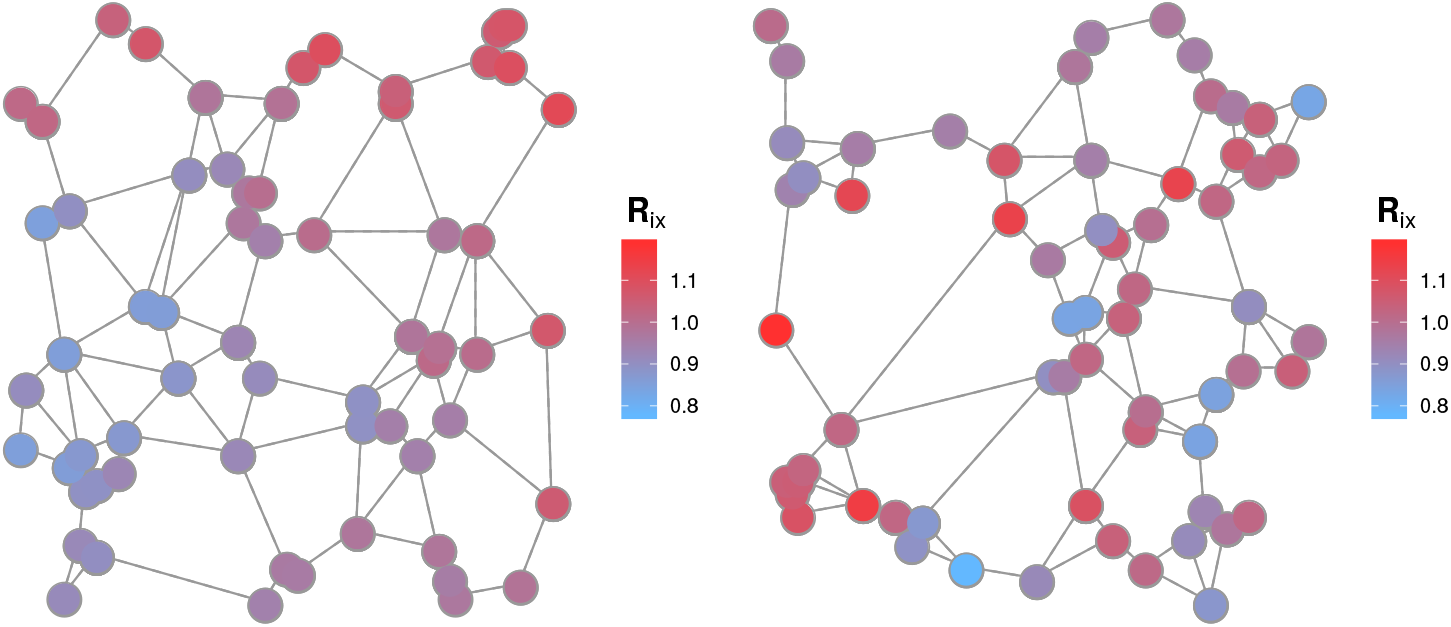
Spatially autocorrelated growth rate distributions. Instrinsic growth rates are sampled from spatially autocorrelated random fields of autocorrelation length *φ* and variance *σ*^2^. Two example distributions are shown, both of *N* = 64, *σ*^2^ = 0.01, with *φ* = 10 (left) and *φ* = 1 (right). See Materials and Methods for details.

**Figure S3.**
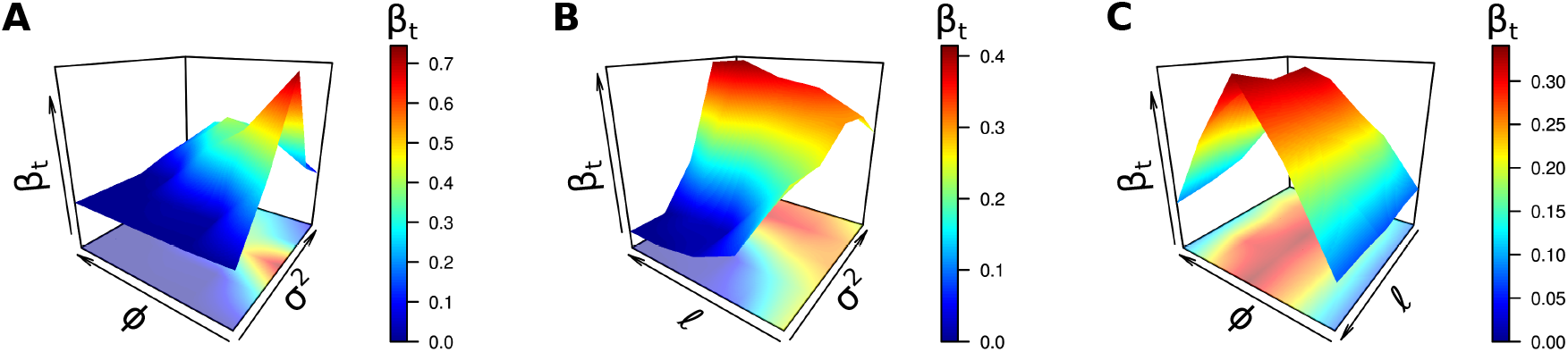
Temporal turnover throughout the spatial parameter space. Temporal *β*-diversity *β_t_* was computed as the mean BC dissimilarity between time points in a time series of 1000 unit times, observed in metacommunities of *N* = 64 patches. Correlation length *φ* was varied in the range 1 to 100, environmental variability *σ*^2^ and dispersal length *ℓ* in the range 10^−2^ to 1, with each parameter combination replicated 10 times. The values of *φ, σ*^2^ and *ℓ* were each plotted on logarithmic axes. In **A** we fixed *ℓ* at 0.5; in **B** *φ* at 10; and in **C** *σ*^2^ at 1.0. See Supplementary Text for details.

**Figure S4.**
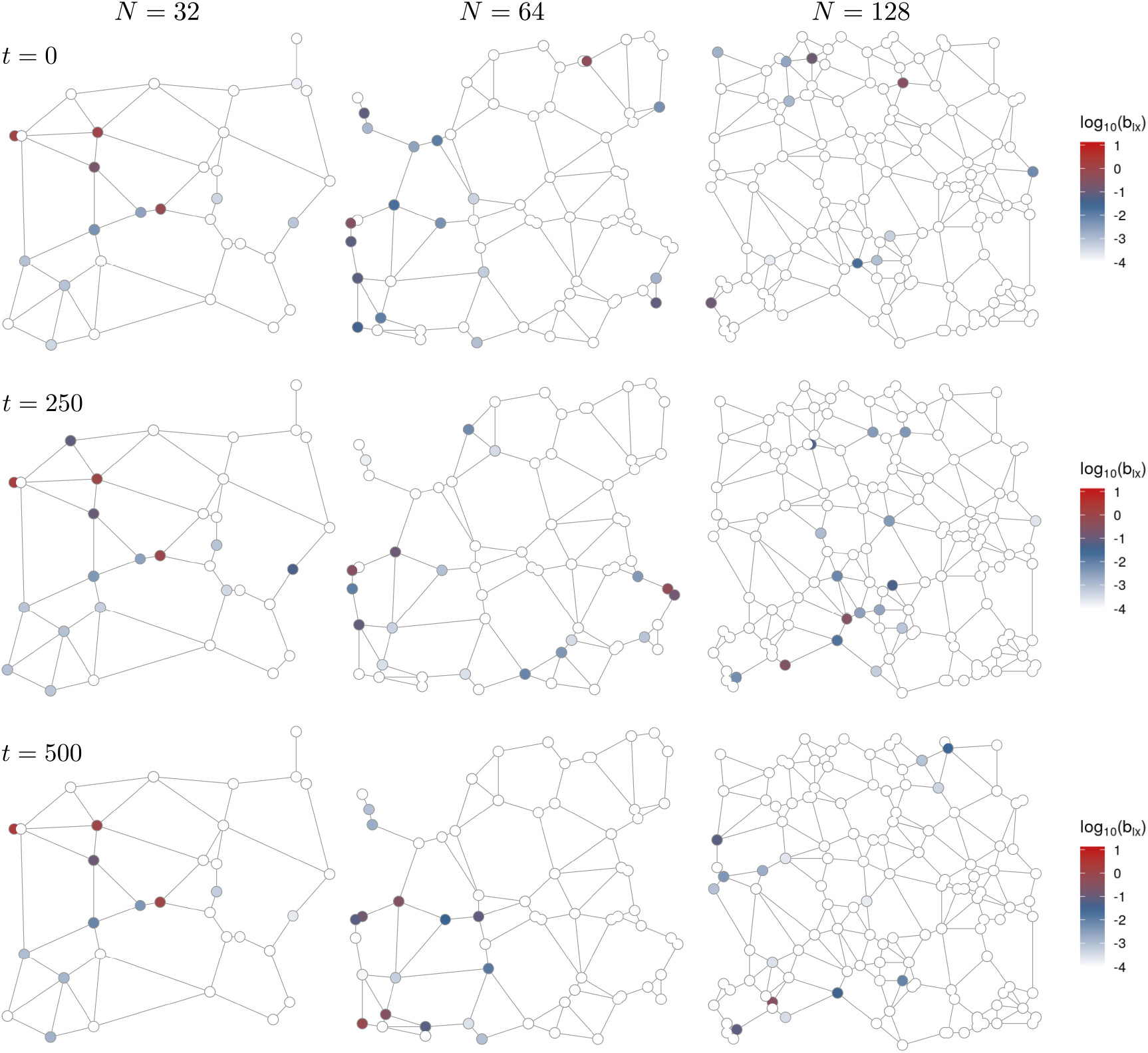
Autonomous metapopulation dynamics in large metacommunity models. In species rich metacommunities of *N* > 8 patches, local biomasses autonomously fluctuate and the variability of those fluctuations increases with metacommunity size. Here we show the instantaneous biomass distributions for a single species in metacommunities of *N* = 32, 64 and 128, at three time points in logarithmic biomass units. For *N* = 32, autonomous fluctuations are largely restricted to the outer extremes of the species’ distribution, while the core range (left of network) remains largely static. For *N* = 64, some patches or regions may be permanently occupied by the focal species, however even in this core range biomass can fluctuate by orders of magnitude. With the emergence of Gleasonian turnover in the high *N* limit no or few patches are permanently occupied and local community composition is no longer well characterized by the core-transient distinction^55,57,60^, which decomposes local communities into populations that are present almost all the time, and those observed only rarely. Hence, for *N* = 128 no obvious core range exists. Note that spatial networks are not shown to scale, the area of the model landscape is ≈N in all cases. *A_ij_* = 0.5 with probability 0.5, *φ* = 10, *σ*^2^ = 0.01, *ℓ* = 0.5. See Main Text for details.

**Figure S5.**
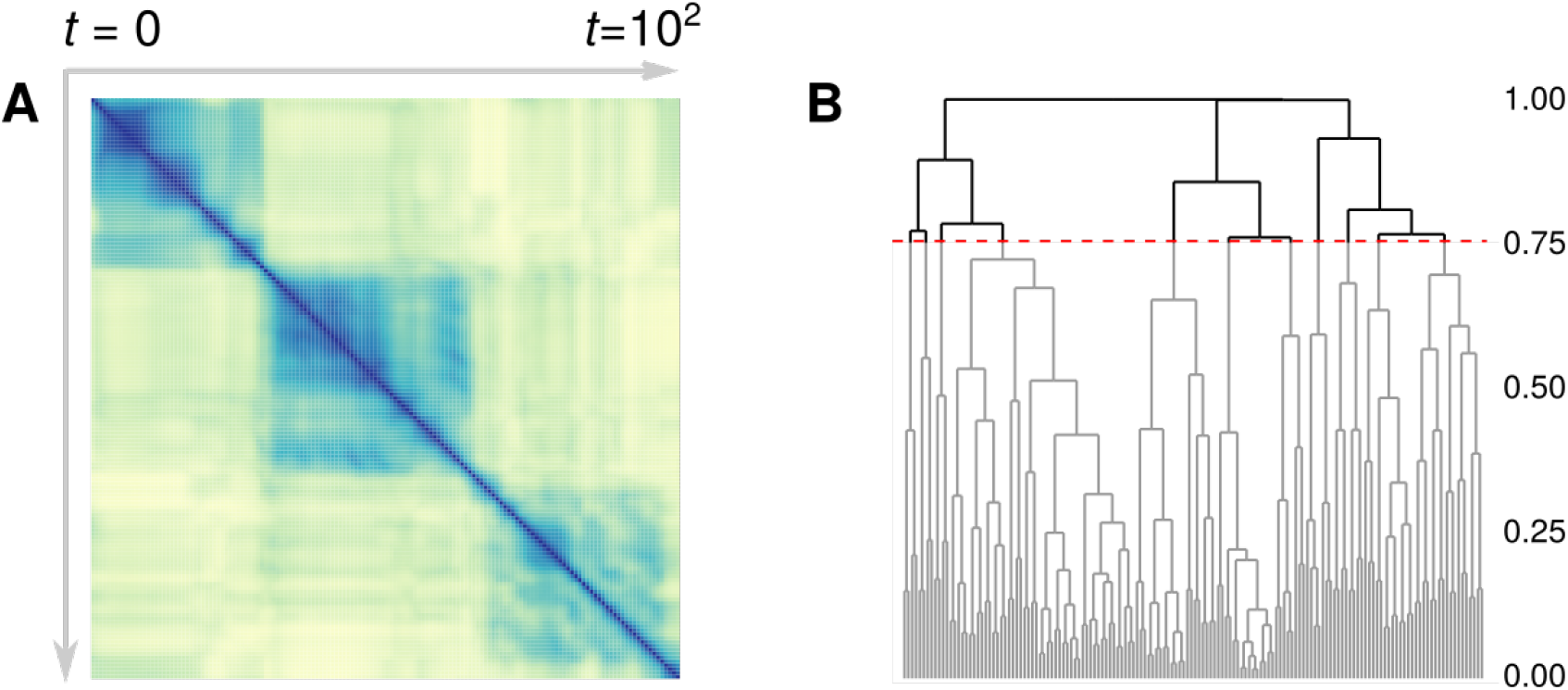
The number of compositional clusters in a community time-series analysed using hierarchical clustering. **A**: Temporal clustering in *local* community composition represented by the block structure of the BC similarity matrix (*N* = 64, 200 unit times shown). **B**: Using hierarchical cluster analysis we approximately quantifies the number of clusters in communit state using a similarity threshold of 75% (red dashed line).

**Figure S6.**
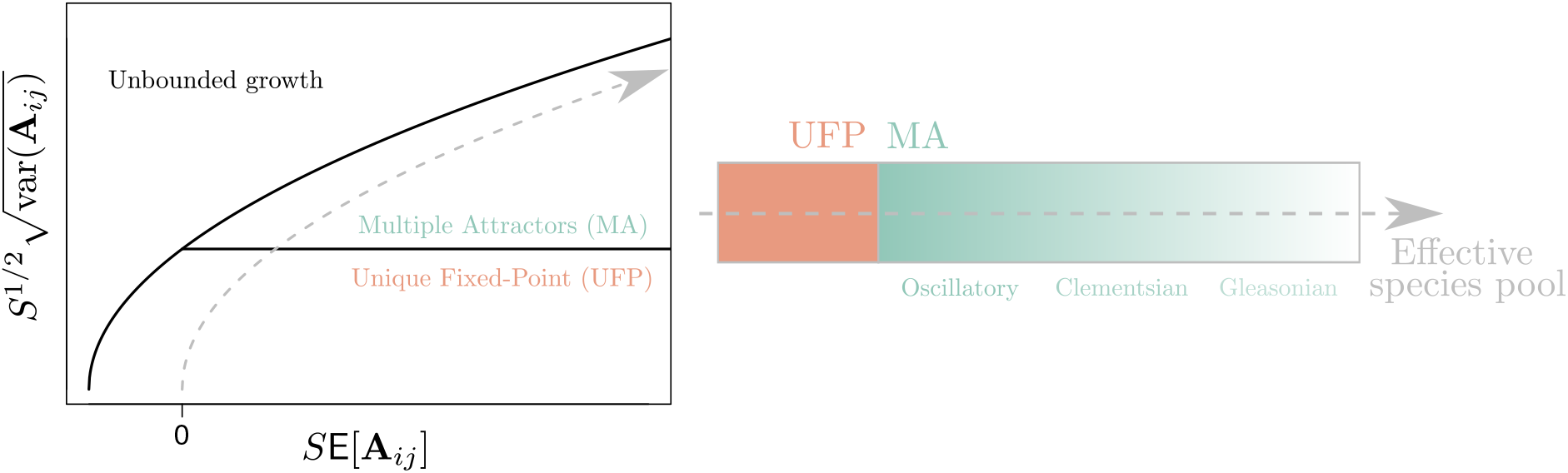
The sharp transition between UFP and MA phases. Reproduction of the phase diagram derived by Bunin^50^ showing the emergence of MA as the size *S* of the species pool increases. In our case, the first and second moments of the distribution in *A_ij_* were fixed. Community state in phase space therefore follows a square root function with increasing *S*, as indicated by the dashed line. (The “Unbounded growth” phase is hence not relevant for our study.) In spatially explicit metacommunity models we observe the emergence of autonomous turnover which transitions from oscillations to Clementsian and finally Gleasonian turnover. See Supplementary Text for details.

**Figure S7.**
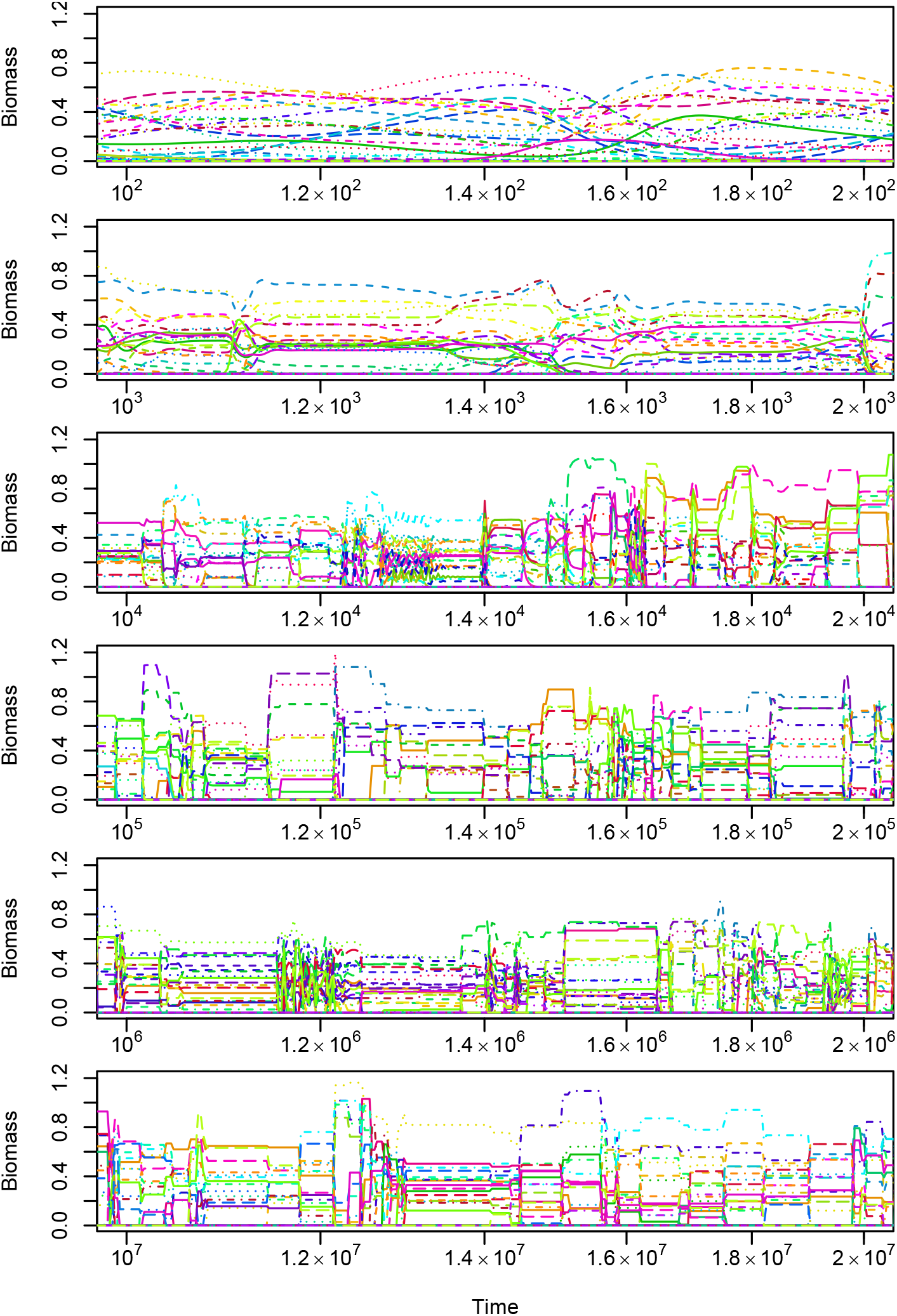
Episodes in the approach of an isolated LV community model to a heteroclinic network. The biomasses of different species are represented by lines of different colours and style. At any moment in time, all but a few of the *S* = 300 species in the system have biomasses close to zero. With increasing simulation times *t* the intervals between the switches in system state, corresponding to transitions from the vicinity of one unstable equilibrium to the next, become longer, while the duration of these transitions remains of the order of magnitude of 10 time units, leading to increasingly sharper transitions on the logarithmic time scale. See Supplementary Text for details.

**Figure S8.**
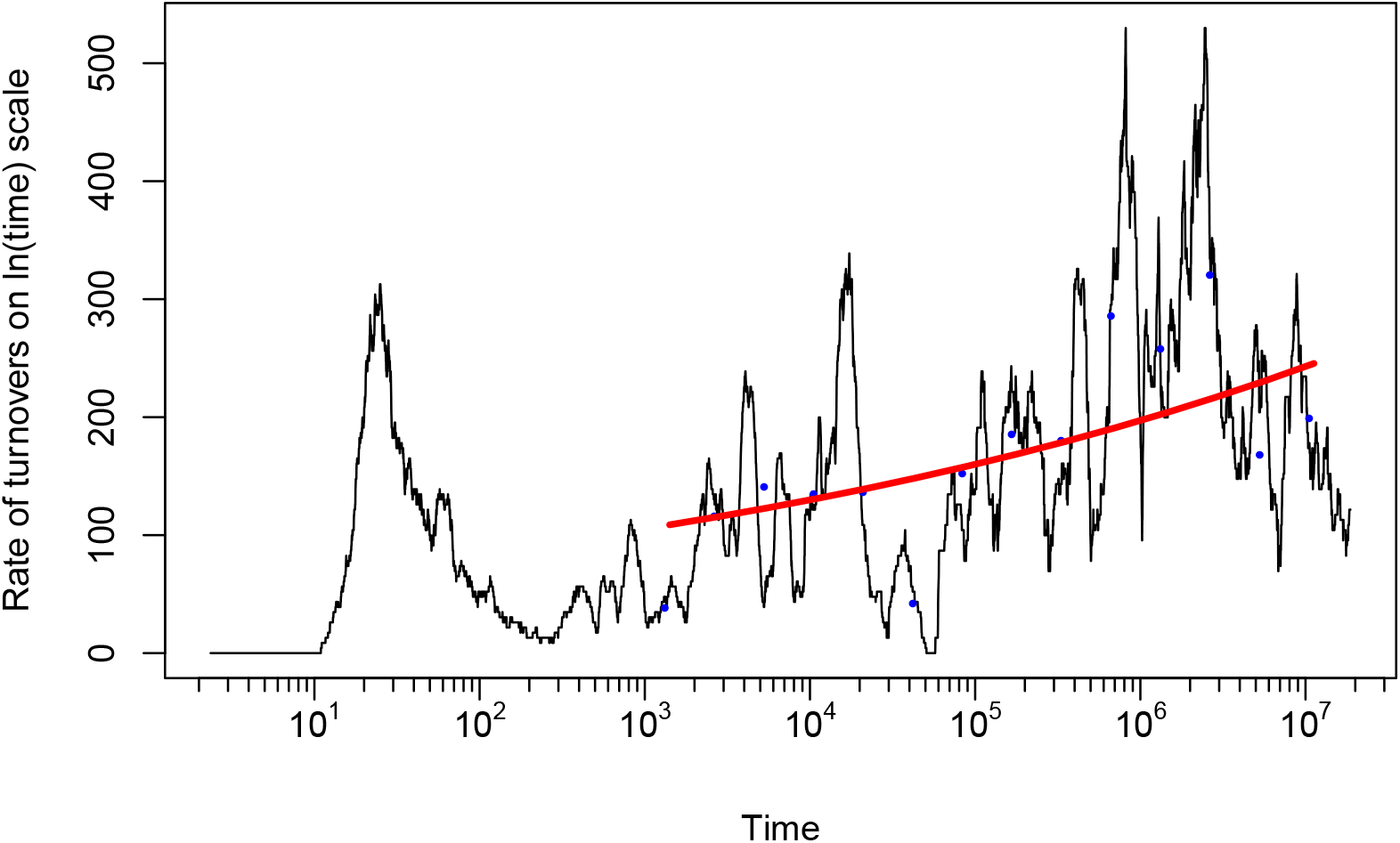
Rate of change in community composition for the simulation shown in Fig. S7. The black line is the moving average over 100 subsequent recordings, blue dots represent averages over non-overlapping adjacent blocks of 300 recordings for *t* ≥ 1000, and the red line a median nonlinear regression of the dots by a power-law (rate) ~ *t^ν^*(*v* = 0.091 ± 0.062, not significantly different from zero). See Supplementary Text for details.

**Figure S9.**
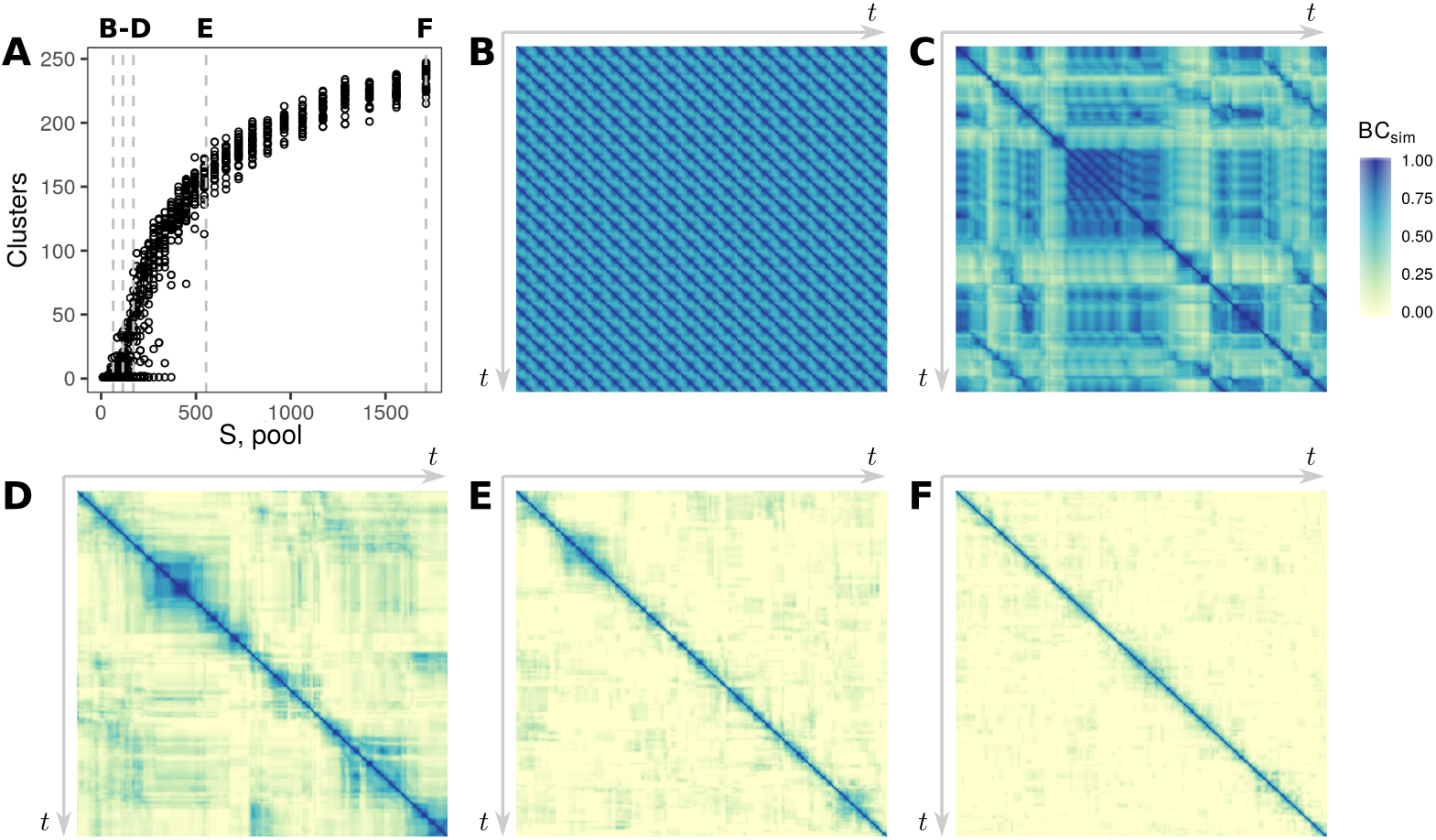
Autonomous turnover in isolated LV communities. **A**: The number of compositional clusters detected as a function of the size of the pool of potential invaders for a propagule pressure, *ϵ*, of 10^−15^ biomass units per unit time. **B-F**: Heatmaps of the pairwise Bray-Curtis similarity for the corresponding time-series (over10^4^ unit times) showing a clear transition from oscillatory to Clementsian turnover and finally to Gleasonian turnover. Dashed lines in **A** show the size of the species pool for which each community time series was generated. *A_ij_* = 0.5 with probability 0.5, *σ*^2^ = 0.01. The parameters *φ* and *ℓ* are not defined for the isolated LV models. See Supplementary Text for details.

**Figure S10.**
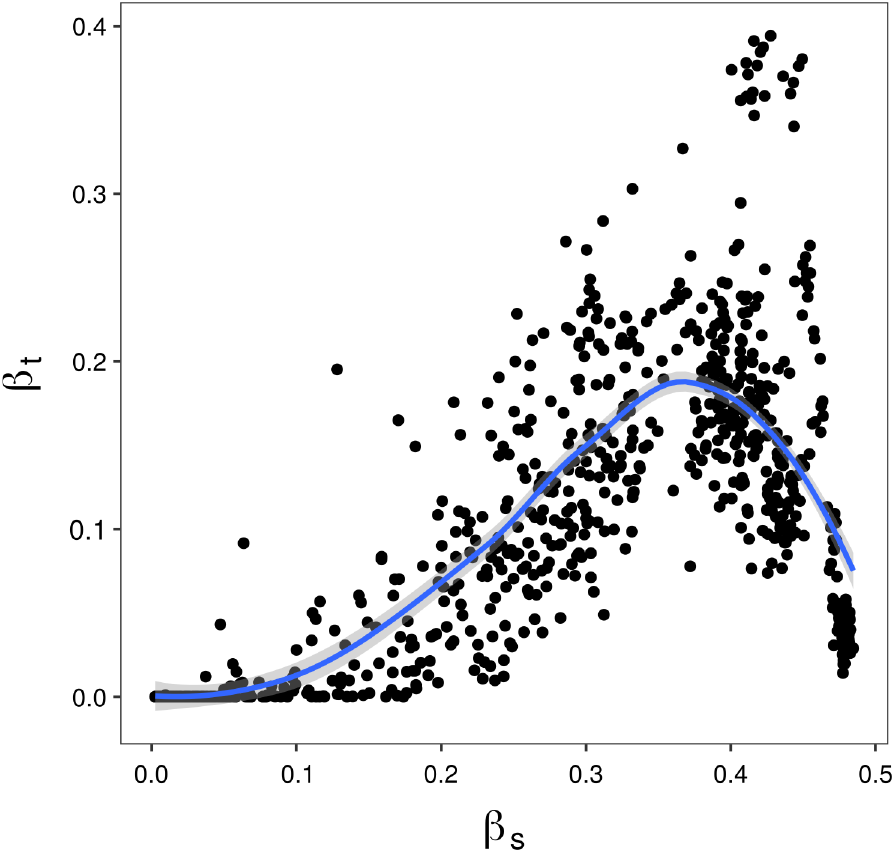
Unimodal relationship between spatial and temporal turnover. Temporal beta diversity, computed during 1000 unit times, plotted against the spatial beta diversity of the local neighbourhood. The number of patches in a local neighbourhood depends on the patch degree, which varies. We therefore use a beta-diversity metric^83^ (based on BC dissimilarity) that is normalises by the number of sites/time-points included in the sub-sample. Both *β*_t_ and *β*_s_ are averages over the metacommunity. The blue line and shaded area represent a locally weighted regression (LOESS smoothing) and 95%C.I.. Parameters *N, φ, σ*^2^ and *ℓ* as in Fig. S3. See Supplementary Text for details.

**Figure S11.**
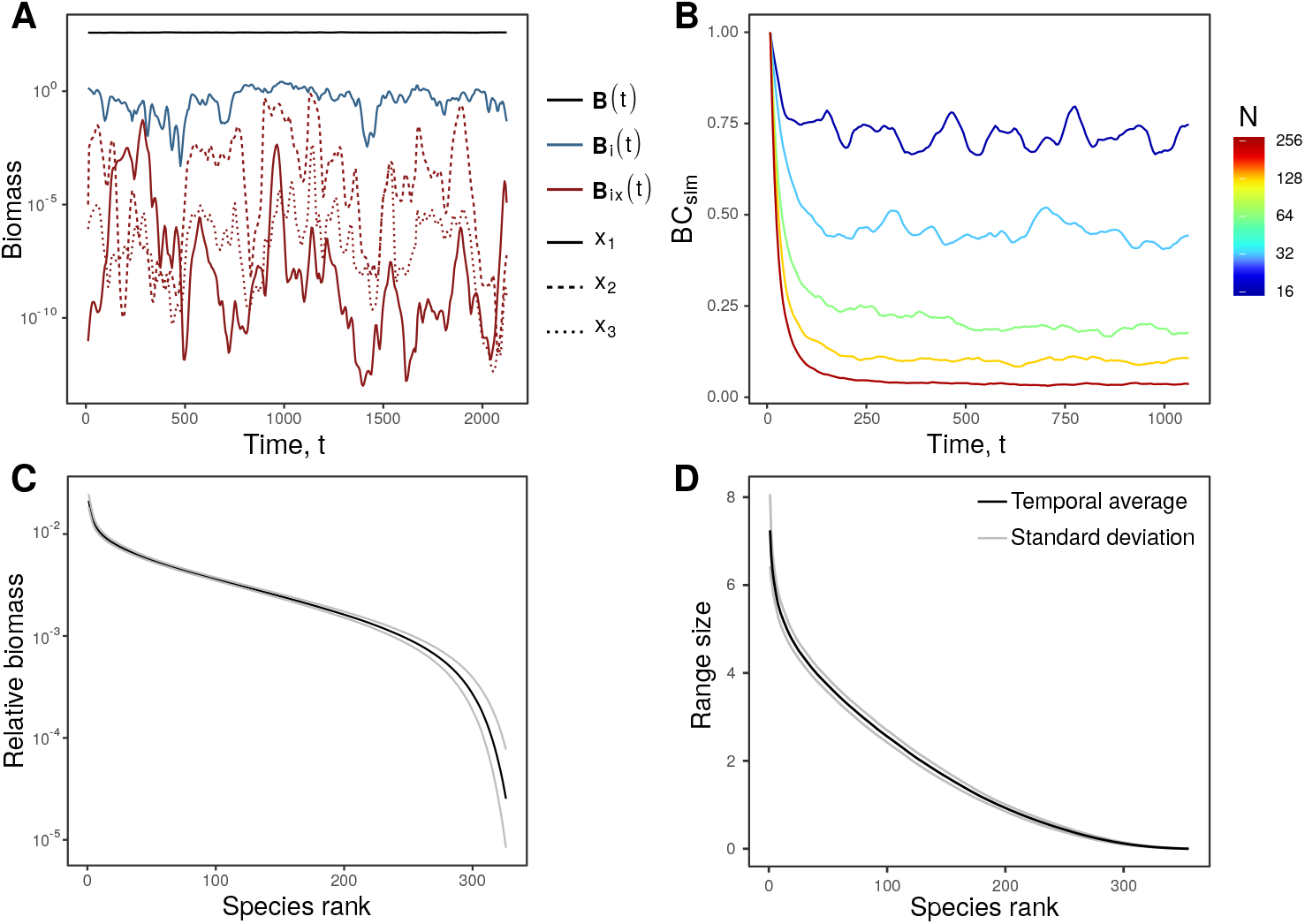
Temporally robust community structure. **A**: We highlight the scale dependence of autonomous population dynamics by showing the biomass of three random local populations of the same species (*B_ix_*, red), of the metapopulation of which they form a part (B_*i*_ = ∑_*x*_ **B**_*ix*_, blue) and finally of the entire metacommunity (B ∑_*i*_∑_*x*_ **B**_*ix*_), black). **B**: Autonomous turnover can be substantial. Here we show the decay of spatially averaged BC similarity from an arbitrary initial composition in metacommunities of *N* = 16, 32, 64, 128, and 256 patches. For large metacommunities undergoing autonomous Gleasonian turnover, the percentage of permanent populations, and hence the temporal BC similarity can drop to zero. **C**: Metacommunity scale relative rank abundance curve, plotted with species ‘identity’ disregarded. The black curve represents the mean biomass observed at a given rank, while grey curves represent the mean ± one standard deviation. This figure highlights the temporally invariant diversity structure at the metacommunity scale. **D**: The temporally averaged rank range size curve, plotted as in C. *A_ij_* = 0.5 with probability 0.5, *φ* = 10, *σ*^2^ = 0.01, *ℓ* = 0.5. *N* = 64 for **A**, **C** and **D**. See Supplementary Text for details.

**Figure S12.**
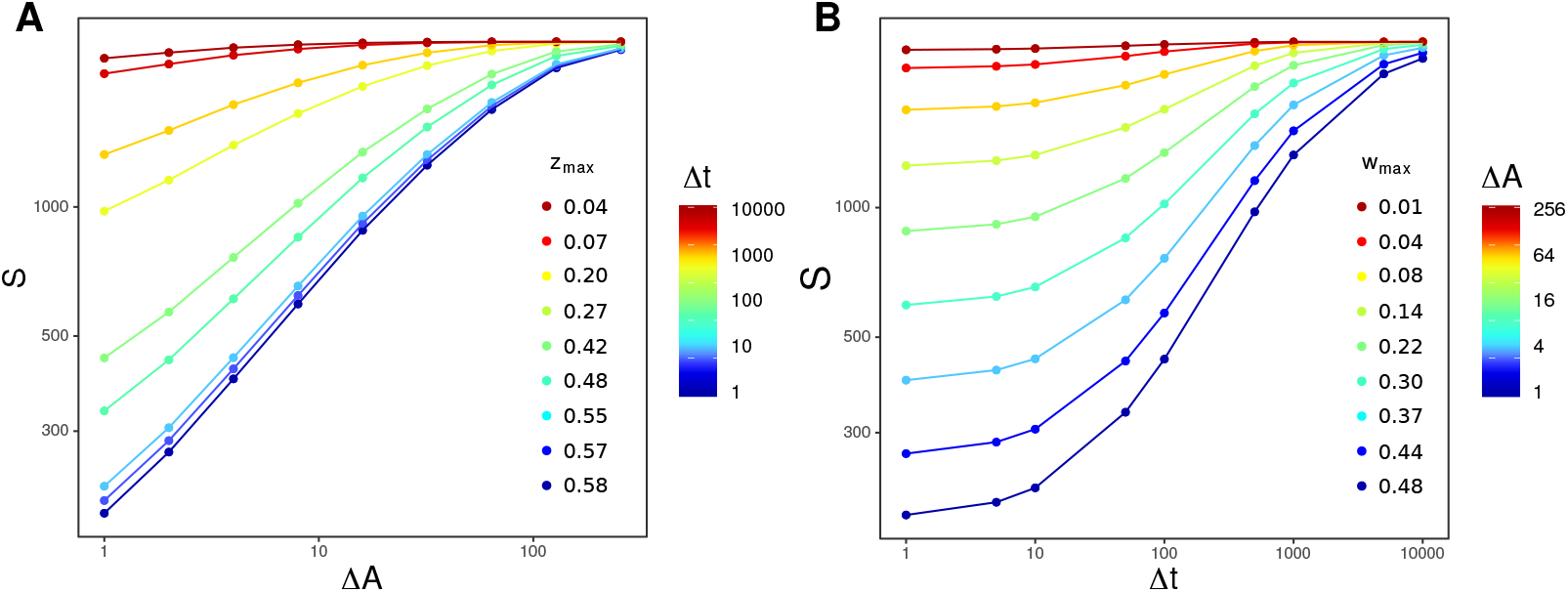
The Species-Time-Area-Relation. The nested SAR (**A**) and STR (**B**) generated using a sliding window approach for a single metacommunity model of *N* = 256. Metacommunity models are closed systems and as such, both the SAR and STR saturate for the large sub-samples. As such we defined the exponents of the STAR by the maximum slopes observed on double logarithmic axes. *A_ij_* = 0.5 with probability 0.5, *φ* = 10, *σ*^2^ = 0.01, *ℓ* = 0.5. See Supplementary Text for details.

## Code availability

Access to custom code used to generate the data analysed in this study is available from the corresponding upon reasonable request.

## Notes

### Competing Interest Statement

The authors have declared no competing interest.

